# Screening by deep sequencing reveals mediators of microRNA tailing in *C. elegans*

**DOI:** 10.1101/2021.01.11.426275

**Authors:** Karl-Frédéric Vieux, Katherine Prothro, Leanne H. Kelley, Eleanor M. Maine, Isana Veksler-Lublinsky, Katherine McJunkin

**Author notes:** The authors wish it to be known that, in their opinion, the first two authors should be regarded as joint First Authors.

## Abstract

microRNAs are frequently modified by addition of untemplated nucleotides to the 3’ end, but the role of this tailing is often unclear. Here we characterize the prevalence and functional consequences of microRNA tailing *in vivo*, using *C. elegans.* MicroRNA tailing in *C. elegans* consists mostly of mono-uridylation of mature microRNA species, with rarer mono-adenylation which is likely added to microRNA precursors. Through a targeted RNAi screen, we discover that the TUT4/TUT7 gene family member CID-1/CDE-1/PUP-1 is required for uridylation, whereas the GLD2 gene family member F31C3.2 – here named <u>G</u>LD-2-<u>r</u>elated <u>2</u> (GLDR-2) – is required for adenylation. Thus, the TUT4/TUT7 and GLD2 gene families have broadly conserved roles in miRNA modification. We specifically examine the role of tailing in microRNA turnover. We determine half-lives of microRNAs after acute inactivation of microRNA biogenesis, revealing that half-lives are generally long (median=20.7h), as observed in other systems. Although we observe that the proportion of tailed species increases over time after biogenesis, disrupting tailing does not alter microRNA decay. Thus, tailing is not a global regulator of decay in *C. elegans*. Nonetheless, by identifying the responsible enzymes, this study lays the groundwork to explore whether tailing plays more specialized context- or miRNA-specific regulatory roles.

## Introduction

microRNAs (miRNAs) are small non-coding RNAs that bind Argonaute (Ago) proteins to form the miRNA-induced silencing complex (miRISC), which represses complementary target mRNAs (1). miRNA-target interactions are important in sculpting gene regulation for normal physiology (2).

Canonical miRNAs are derived from longer primary transcripts (pri-miRNAs) from which a hairpin-like secondary structure is recognized and excised by the Microprocessor complex, made up of the ribonuclease Drosha and its protein cofactor DGCR8/Pasha (3). This excised hairpin, known as the miRNA precursor (or pre-miRNA), is exported to the cytoplasm where its loop is cleaved by Dicer (3). One strand of the resulting miRNA duplex (the “guide” strand) is then preferentially loaded into Ago, and the passenger strand (“star” strand) is degraded (3).

In contrast to biogenesis, the process of miRNA turnover is poorly understood, despite turnover’s equivalent importance in determining miRNA abundance. In general, miRNA half-lives are long, on the order of ~20h, with some miRNAs exhibiting characteristically shorter half-lives (4–8). Certain physiological signals and developmental contexts can accelerate miRNA turnover by poorly understood mechanisms, such as neuronal activity which globally destabilizes miRNAs in the retina (9).

Untemplated nucleotide additions to the 3’ end (“tailing”) can regulate both biogenesis and decay of miRNAs through multiple mechanisms. These modifications are carried out by a class of enzymes known as terminal nucleotidyl transferases (TENTs), which also have other roles in regulating mRNAs and viral genomes (10, 11).

The most well-understood examples of tailing regulating miRNAs involve uridylation of miRNA precursors. This type of tailing generally impacts atypical precursors that lack the canonical 2-nt single-stranded 3’ overhang which is generated by Microprocessor cleavage and recognized by Dicer (12). The type of uridylation and its consequence depends on the length of the 3’ overhang, where 1-nt overhangs are mono-uridylated to favor biogenesis and further recessed ends are oligo-uridylated resulting in decay of the precursor (12). Precursor uridylation can also regulate biogenesis by shifting the site of Dicer cleavage, which can result in alternative miRNA guide strand selection (arm-switching) (13). Oligo-uridylation and decay of the *let*-7 precursor are induced even in the context of a canonical 2-nt 3’ overhang by binding of the RNA-binding protein LIN-28 (14, 15). Finally, non-canonical Microprocessor-independent miRNA precursors derived from short introns (mirtron precursors) are also highly uridylated, inhibiting their biogenesis in *Drosophila* (16, 17).

Tailing can also occur on mature miRNAs, again with a wide variety of outcomes. During embryonic development of *Drosophila*, fish, and mice, terminal adenylation of maternally-contributed miRNAs induces sequence-non-specific miRNA decay at the maternal to zygotic transition (18). In contrast to the clearance of adenylated maternally-contributed miRNAs, terminal adenylation of mature miRNAs is stabilizing and activating for miR-122 and a subset of other miRNAs (19–21).

Through a process known as target-directed miRNA degradation (TDMD), target RNAs with extensive complementarity cause the 3’ end of miRNAs to be tailed and trimmed, concomitant with miRNA destabilization (22–31). While tailing was initially assumed to be causative of miRNA decay in this context, recent studies demonstrate examples in which tailing is dispensable for TDMD, suggesting that 3’ tailing may be a symptom of miRNA 3’ end display rather than an upstream regulatory event in TDMD (31–35).

Beyond effects on stability, 3’ miRNA tailing can even alter target repertoire (36). Overall, tailing of miRNA precursors and mature miRNAs has a wide range of regulatory consequences. How tailing is coupled to these various outcomes in a miRNA- and context-dependent manner is not understood.

In this work, we set out to examine the role of tailing in miRNA turnover in a model organism *in vivo.* We first use deep sequencing and meta-analysis of published data to characterize miRNA tailing in *C. elegans*. We perform a candidate RNAi screen by deep sequencing to discover the enzymes responsible for tailing, demonstrating that <u>t</u>erminal <u>u</u>ridylyl<u>t</u>ransferase 4 (TUT4)/TUT7 ortholog <u>c</u>affeine <u>i</u>nduced <u>d</u>eath homolog 1 (CID-1) is required for U-tailing, and defective in <u>g</u>erm <u>l</u>ine <u>d</u>evelopment 2 (GLD2) ortholog F31C3.2 – here named <u>GLD</u>-2-<u>r</u>elated <u>2</u> (GLDR-2) – is responsible for A-tailing. Finally, we use acute inactivation of miRNA biogenesis to measure miRNA turnover and show that, while U-tailing increases as miRNAs age, this is correlative not causative of miRNA turnover. Overall, this work importantly defines ancestral functions of TENT families that modify miRNAs, and – by defining the required enzymes – lays the groundwork for future studies to determine whether miRNA tailing in *C. elegans* may play context-dependent or miRNA-specific functions.

## Methods

### *C. elegans* Maintenance and RNAi

N2 (wild type) and SX1137 (*pash*-*1(mj100)* I) worms were maintained at 15°C. For RNA samples, L1s were synchronized by alkaline hypochlorite lysis of gravid adults followed by hatching in M9 with 1mM cholesterol at 15°C for 48-72h. L1s were plated on NGM seeded with either OP50 or RNAi bacteria and maintained at 15°C until early day one of gravid adulthood (until just a few embryos were laid on the plate, ~76h after plating). (This time point was chosen because it achieved the best synchrony of SX1137, which tends to develop slightly asynchronously, even at permissive temperature.) This time point was harvested for N2 samples, SX1137 samples in the RNAi screen, and served as 0h in time courses involving SX1137. Additional SX1137 plates were shifted to 25°C, and samples were collected at the indicated time points after upshift for time courses on OP50 or RNAi. *pup*-*1(tm1021)*, *pup*-*2(tm4344)*, and *pup*-*1/*-*2(om129)* mutations were maintained over a balancer chromosome at 15-20°C; RNA was isolated from homozygous F2 (M-Z-) 1-day adult hermaphrodites raised at 22°C.

RNAi plates were supplemented with 1μg/ml IPTG and 1μg/ml Carbenicillin (and poured no more than two weeks prior to use and stored at 4°C). The RNAi bacteria (HT115 expressing an RNAi vector) were prepared from single clones derived from the Ahringer library (Source Bioscience), the Vidal library (Source Bioscience), or custom RNAi vectors (Table S1). All previously-reported RNAi phenotypes were observed. Furthermore, a robust RNAi response in each sample was confirmed by the presence of abundant siRNAs mapping to the targeted locus.

### Small RNA Sequencing

Total RNA was isolated by Trizol extraction. Spike-ins were added at a final concentration of 0.1pg/μl (OP50 samples) or 1pg/μl (RNAi samples) in 100ng/μl total RNA. See Table S2 for spike-in information. Libraries were prepared using the NEBNext Small RNA Library Prep set for Illumina with the following modifications. The first size selection was performed after the RT reaction by excising 65-75nt cDNA from an 8% denaturing acrylamide gel. PCR products were again size selected using a Pippin prep or 6% native acrylamide gel. Input RNA and final library sample quality were assessed using a Bioanalyzer. Equimolar amounts of each library were pooled; up to 24 indexed samples were pooled together for sequencing. For *pup*-*1(tm1021)*, *pup*-*2(tm4344)*, and *pup*-*1/*-*2(om129)* libraries, ~18-35 bp inserts were size-selected. Sequencing was performed on a NextSeq500 with a minimum of 10M single-end reads of minimum length 50 nt.

### Data analysis

Deep sequencing data were analyzed using the miTRATA web interface (37) after preprocessing the data using miTRATA’s accompanying Python3-based preprocess.seq pipeline, installed on the NIH High Performance Computing Cluster in a dedicated miniconda environment. The mature *C. elegans* miRNAs from miRBase Release 22.1 (38) were used for miTRATA mapping. A custom C++ script reformatted the miTRATA output by consolidating read numbers for all reads meeting the following criteria into a tabular format: a tail:head ratio > 0.12 (16) and a tail length ≤ 3nt. These reformatted tailing data were further analyzed in RStudio. (One non-unique miRNA pair was of sufficient abundance to be included in tailing analysis – *mir*-*44/45*-*3p*; *mir*-*45*-*3p* was therefore culled from tabular data to avoid plotting this data twice.) To calculate reads per million, reads were normalized by the number of genome-mapping reads in the library (Table S3). The number of genome-mapping reads was determined using Bowtie (39) to align to the *C. elegans* genome (WS215), with the arguments -v 3 -f -B 1 -a --best --strata. Alignments were then filtered based on the length of the reads and the number of mismatches as follows: for sequence lengths 16-17, 18-19, 20-24, or >24: zero, one, two, or three mismatches were allowed, respectively. Reads passing this threshold were considered “genome-mapping reads”. A custom bash/R pipeline was used to calculate trimming relative to canonical miRNA length from the miTRATA output files.

### Statistical Analysis

Fisher’s exact test for enrichment of tailing on 3p- or 5p-derived miRNAs was performed by counting miRNAs as above or below 1% tailed, and pooling these counts from three biological replicates to generate the contingency table, which was analyzed in GraphPad QuickCalcs. For the RNAi screen, GraphPad Prism v8.4.3 was used to perform one-way repeated measures ANOVA followed by Dunnett’s multiple comparison test. All RNAi conditions were compared to an empty vector sample, and this comparison was repeated for each empty vector sample. RNAi conditions that were significant in comparison to each empty vector sample are highlighted, with the least significant p-value reported. All miRNAs >50 RPM in all empty vector replicates were analyzed. To determine miRNA half-lives, for each miRNA with >50 RPM at 0h in both replicates, fold change at each time point relative to 0h was calculated using spike-in-normalized reads. GraphPad Prism v8.4.3 was used to fit a one phase decay non-linear regression model with the following constraints: Y0=1, Plateau=0, K>0.

### Phylogenetic analysis

MEGA-X (40) was used to build phylogenetic trees of orthologous genes (to CID-1 and PUP-2, or to GLD-2 and F31C3.2) identified in WormBase ParaSite version WBPS15 (41). Orthologs from select non-nematode species were included as shown (Figures S7 and S9). The classification of nematode species into clades was downloaded from the same WormBase Parasite version WBPS15. Additional trees containing only *C. elegans* genes and common model organisms are shown in Figures 2 and 3. All alignments were generated using Clustal W (Multiple Alignment Gap Opening Penalty=10, Gap Extension Penalty=0.2, Negative Matrix Off, Delay Divergent Cutoff 30%). Maximum Likelihood method was used in MEGA-X, with no test of phylogeny, JTT model of substitution, uniform rates among sites, no subsetting (“use all sites”), NNI, automatic initial tree, and no branch swap filter.

### Genome editing

To generate the F31C3.2/*gldr*-*2(cdb187)* mutant, adult N2 hermaphrodites were injected with Cas9/gRNA RNPs to perform CRISPR. The gRNAs (gKFV05 and gKFV06) were designed to cleave within 300nt upstream and downstream of the F31C3.2/*gldr-2* ORF and ordered from IDT as Alt-R crRNAs. Both guides were used at 2μM in the injection mix. 500ng/μl of oKFV101 (Table S2) was used as a ssDNA oligo repair donor to produce a null deletion mutant. The final injection mix also included 2μM IDT Cas9 as well as 2μM of a *dpy*-*10* gRNA and 820nM of a *dpy-10* ssDNA oligo repair donor to generate a Dpy and Rol selection marker (42). See Supplemental Table S2 for gRNA, donor, and genotyping primer sequence details.

### Transgenesis and TDMD sample preparation

Constructs for TDMD triggers were generated using pCFJ420 (*myo*-*3p*::*gfp*::*h2b*::*tbb-2* terminator) as a backbone (43). For each trigger, sequences with high complementarity to the miRNAs of interest were ordered as complementary oligonucleotides, annealed at 12.5μM each in IDT duplex buffer, digested with BsmI, and cloned into the *tbb*-*2* 3’-UTR region of pCFJ420 (also digested with BsmI). See Supplemental Table S2 and Figure S15 for sequence details. The resulting plasmids (pKFV18, PKFV19, and pKFV20) were then injected into adult N2 hermaphrodite gonads to generate extra-chromosomal arrays. Injection mixes included pKFV18, pKFV19, pKFV20, or pCFJ420 (no-binding site negative control) along with the following selection markers: pDD282 (for Hygromycin resistance), pRF4 (for Rol phenotype) and mCherry expression plasmids (pGH8, pCFJC90, and pCFJ104) (44, 45). See Table S2 for injection mix details. Two to three stably transmitting arrays were isolated for each injected construct. To generate samples for deep sequencing, mixed populations were bleached in sodium hypochlorite and synchronized overnight in M9 supplemented with 0.3mg/mL Hygromycin to select for worms carrying the extra-chromosomal array, then grown at 20°C on standard NGM seeded with OP50 until early day one of gravid adulthood. miRNA-Taqman assays (ThermoFisher Scientific) were performed according to manufacturer’s instructions, using synthetic miRNAs to generate a standard curve for absolute quantification. Digital droplet PCR (ddPCR) was performed as a two-step RT-qPCR by first synthesizing oligo(dT) primed cDNA using the iScript™ Select cDNA Synthesis Kit (Biorad), then running ddPCR via the standard EvaGreen program on a Biorad QX200 ddPCR system and analyzing in Quantasoft software (see Table S2 for primer sequences).

## Results

### Patterns of miRNA tailing in *C. elegans* suggest different substrates than in flies and vertebrates

To determine the nature and prevalence of miRNA tailing in *C. elegans*, we first cloned and deep sequenced small RNAs from wild type animals. To ensure reliable measurements of tailing, only miRNAs with greater than 50 reads per million were included in tailing analysis. To determine the level of background in our sequencing and computational pipeline, ten synthetic miRNAs (whose sequences are endogenous to plants and not present in *C. elegans* – Table S2) were spiked in to purified total RNA before preparation for deep sequencing. Although these RNA species were never present in the context of cellular lysate, up to 1% tailing was detected; therefore, tailing levels measured below 1% (delineated by dashed line on all tailing plots) are low confidence and could arise due to PCR or sequencing errors.

Most tailing consisted of a single nucleotide addition (Figure 1A). Tails of two or more nucleotides were much rarer, and so we focused our analysis on further characterization of the single-nucleotide tails. Three replicates of adult animals showed strong correlation in proportions of tailing (Figure S1). Modest levels of miRNA tailing were observed, and tailing in *C. elegans* consisted primarily of uridylation with a much lower frequency of adenylation (Figure 1B, Table S4). This is in contrast to humans and flies, where both adenylation and uridylation are prevalent (46–49). In addition, a single miRNA, *mir*-*83*-*3p*, was highly cytidylated in all replicates (Figure 1B, Figure S1), a modification that is rarely observed in animals, but was recently reported as a frequent addition to plant miRNA precursors (50).

**Figure 1.**
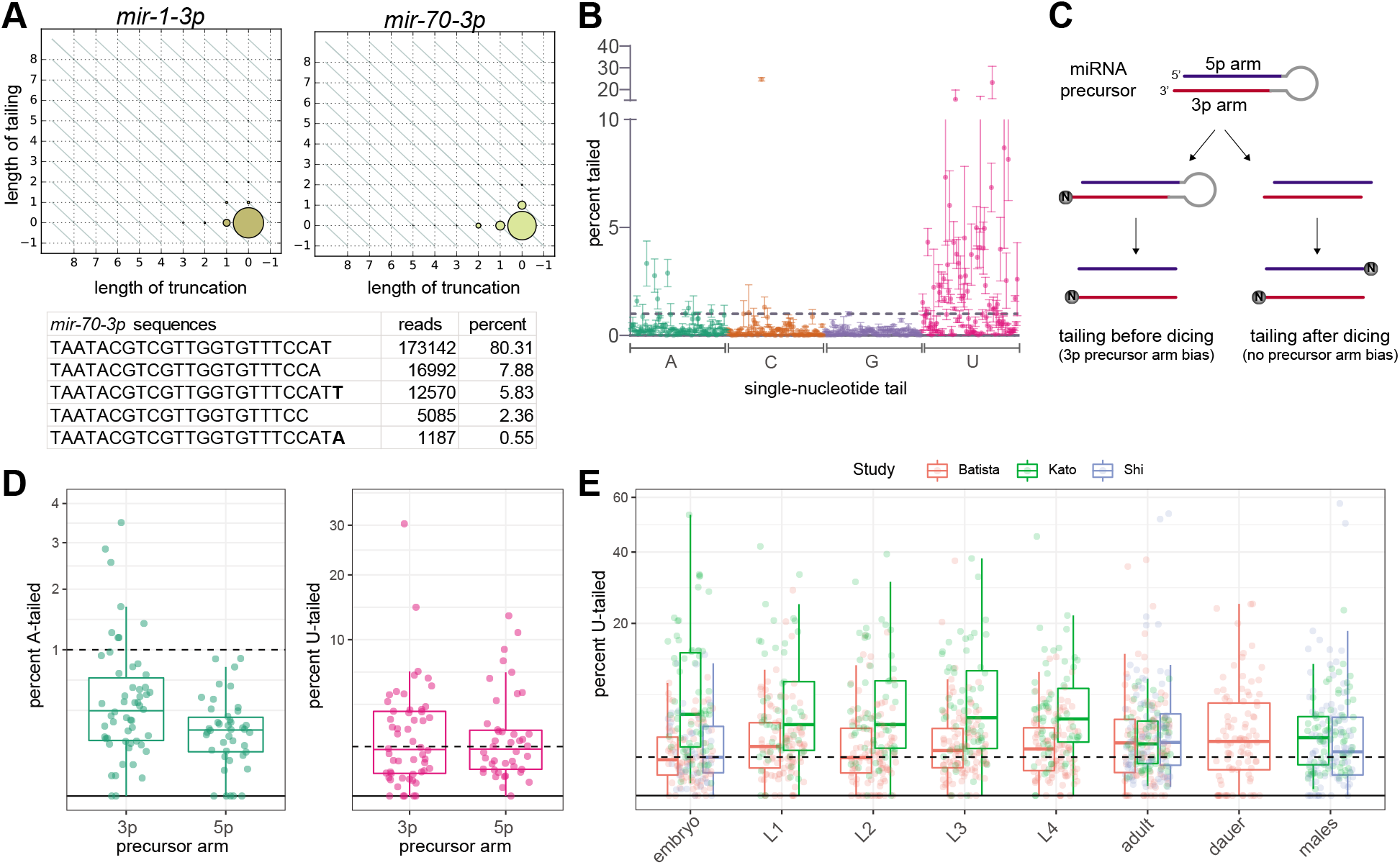
Characterization of miRNA tailing in *C. elegans*. (A) 2D matrix representing the truncation and tailing status of the indicated miRNAs. The sizes of the dots represent the proportion of reads, with the canonical sequence at zero on both axes, and increasingly trimmed or tailed species plotted to the left on the x-axis or upward on the y-axis, respectively. Bottom: Number of reads of most abundant *mir*-*70*-*3p* species in one wild type adult library, with the canonical sequence being the most abundant. (B) Mean and standard deviation of prevalence of single nucleotide 3’ terminal additions of each indicated nucleotide from three biological replicates. (C) Schematic of miRNA tailing preceding or following Dicer-mediated cleavage of the miRNA precursor. (D) Comparison of prevalence of mono-adenylation and mono-uridylation on 3p- versus 5p-derived miRNAs. One biological replicate is shown. (E) Meta-analysis of three published datasets shows tailing across *C. elegans* developmental stages. Prevalence of mono-uridylation of miRNAs across development is shown. Only miRNAs with >50 RPM in the indicated library were analyzed. (B, D) Only miRNAs with > 50 RPM in all biological replicates were analyzed. (B, D-E) Each dot represents an individual miRNA.

**Figure 2.**
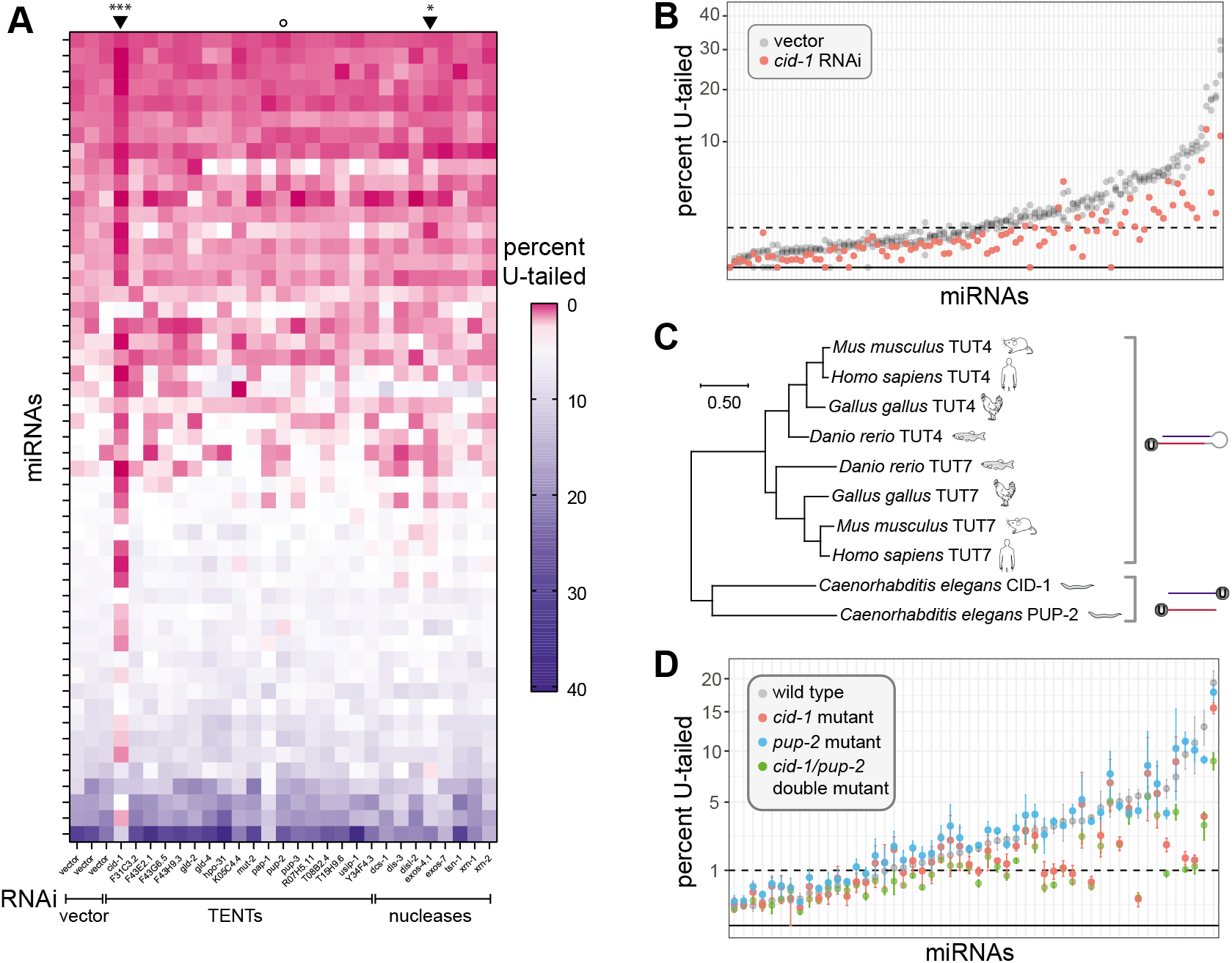
miRNA uridylation requires CID-1. (A) Heatmap summarizing percent of mono-uridylation across each miRNA (rows) in each RNAi condition (columns). All miRNAs with >50 RPM in all empty vector replicates and an average of >1% mono-uridylation across the three empty vector replicates are shown. Arrowhead above heatmap indicates column significantly different than vector (*cid*-*1* and *exos*-*4* RNAi). \*\*\**p*-value = 0.0003, * *p*-value = 0.016. Open circle indicates *pup*-*2* RNAi, which does not have an effect. (B) Percent mono-uridylation in vector or *cid-1* RNAi. Each column is an individual miRNA. All miRNAs with >50 RPM in all empty vector replicates are shown. Three biological replicates of empty vector are shown in gray. (C) Phylogenetic relationship of CID-1, PUP-2, TUT4, and TUT7. (Note the following alternative names for these genes CID-1/CDE-1/PUP-1, PUP-2, TUT4/TENT3A/ZCCHC11/PAPD3, and TUT7/TENT3B/ZCCHC6/PAPD6.) (D) Percent mono-uridylation in indicated genotype. All miRNAs with >50 RPM in all wild type replicates are shown. Mean and standard error are shown for three biological replicates per genotype.

**Figure 3.**
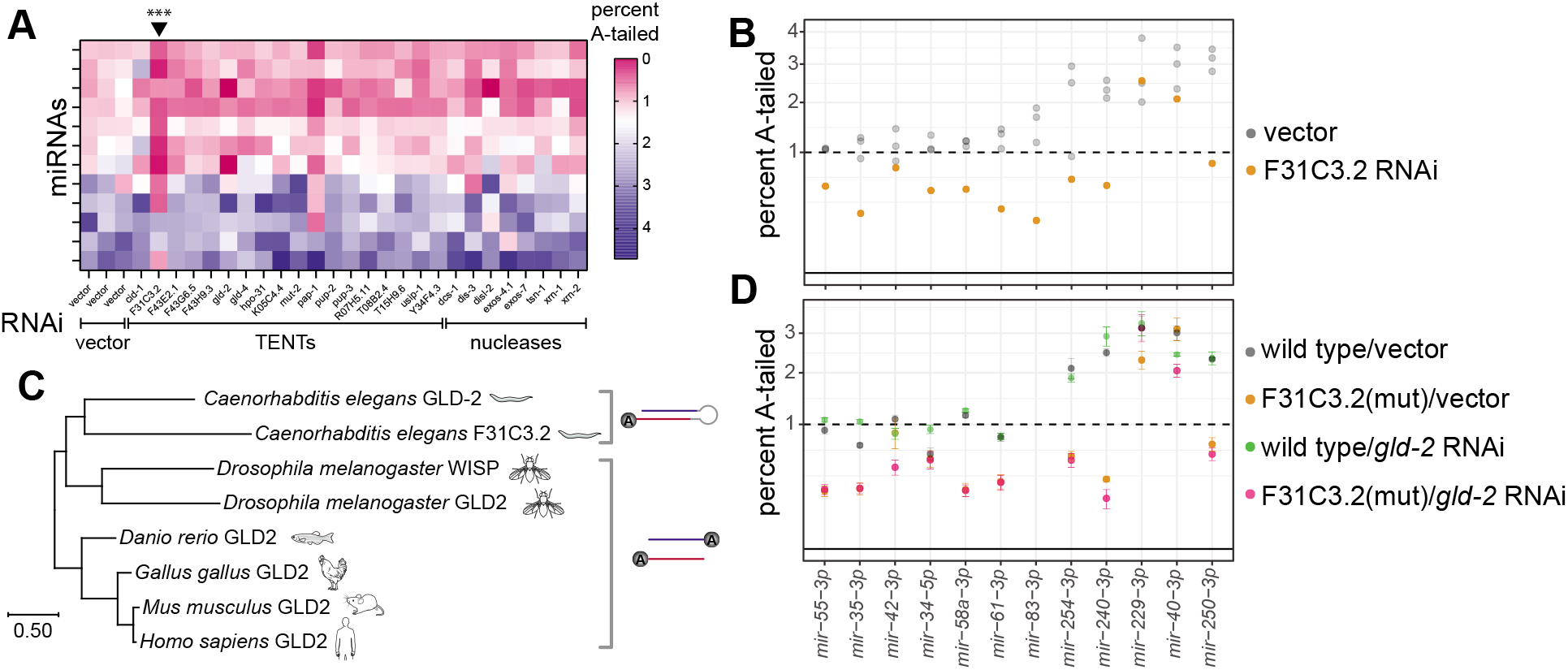
miRNA adenylation requires F31C3.2/GLDR-2. (A) Heatmap summarizing percent of mono-adenylation across each miRNA (rows) in each RNAi condition (columns). Arrowheads above heatmap indicate columns significantly different from vector, corresponding to F31C3.2 RNAi, which reduces adenylation. \*\*\**p*-value = 0.0003 (B) Percent mono-adenylation in vector or F31C3.2/GLDR-2 RNAi. Three biological replicates of empty vector are shown in gray. (A-B) All miRNAs with >50 RPM in all empty vector replicates and an average of >1% mono-adenylation across the three empty vector replicates are shown. (C) Phylogenetic tree of F31C3.2/GLDR-2, GLD-2, and WISP. (Note that GLD2 in humans is also known as TENT2/TUT2/PAPD4/APD4.) (D) Percent mono-adenylation in indicated genotypes/RNAi. Mean and standard error for four biological replicates are shown. MiRNAs shown are the same as those in (A-B).

Either the mature miRNA species or the miRNA precursor may be the substrate of the observed tailing. If the vast majority of observed tailing events occur on miRNA precursors, then miRNAs derived from the 3’ arm of the precursor hairpin (3p) would be expected to bear more terminal additions than those derived from the 5’ arm of the hairpin (5p) since the 3’ end of the 5p arm is not accessible for tailing prior to Dicer-mediated cleavage (Figure 1C). In vertebrates and *Drosophila*, miRNA precursors are the primary substrate of uridylation, though mature miRNAs can also be uridylated (33, 46–48, 51). In contrast, 3p and 5p miRNAs displayed similar levels of uridylation in *C. elegans* (Figure 1D), indicating that U-tailing likely predominantly occurs on mature miRNA species. Untemplated adenylation generally occurs on mature miRNAs in vertebrates, and a modest 3p bias suggests that both precursor and mature miRNAs are adenylation substrates in *Drosophila* (46–48). In contrast, A-tailing in *C. elegans* shows a strong bias, occurring primarily on 3p-derived miRNAs (Figure 1D); miRNAs displaying above 1% A-tailing were significantly enriched in the 3p-derived class of miRNAs (Fisher’s exact test p-value < 0.0001). This suggests that A-tailing in *C. elegans* occurs predominantly on miRNA precursors.

Because global developmentally-regulated tailing has been demonstrated in other models (18), we examined published datasets from developmental time courses to determine whether tailing of *C. elegans* miRNAs is correlated with any particular developmental stage (52–54). No consistent global differences in rates of uridylation or other types of tailing were observed (Figure 1E and Figure S2A, Table S5). We were especially interested in oocyte and early embryo samples since miRNA adenylation is highly prevalent in these time points and leads to turnover of maternally-contributed miRNAs in other species (18), but neither adenylation nor uridylation was globally dynamic during the course of early embryonic development in *C. elegans* (Figure S2B, Table S5) (55). When examining the most highly tailed miRNAs from our data (Figure 1B), these miRNAs were largely expressed fairly evenly across development, with the exception of the embryo-enriched *mir*-*35*-*42* family of miRNAs (of which one member was among the top twelve U-tailed miRNAs and three were among the top twelve A-tailed miRNAs) (Figure S3). When examining the tailing of the highly tailed miRNAs across development, tailing was approximately equally prevalent across development (Figure S4).

### miRNA uridylation requires CID-1

To determine what role tailing may play in regulating miRNAs in *C. elegans*, we next set out to identify the enzymes responsible for these modifications. To this end, we performed RNAi against a list of candidate terminal nucleotidyl transferases (TENTs). For each knockdown, we performed deep sequencing of small RNAs, and compared the proportion of tailed isoforms to those in three biological replicates of empty vector. Of RNAi against 18 candidate tailing enzymes, only the RNAi targeting *cid*-*1* significantly abrogated miRNA U-tailing (*p*-value = 0.0003, Figure 2A-B, Figure S5, Table S6), and this was reproduced in independent experiments (Figure S6A). Knockdown of *cid*-*1* decreased U-tailing of both 3p- and 5p-derived miRNAs equally, suggesting that CID-1 acts on mature miRNAs (Figure S6B). Though *cid*-*1(lf)* was previously reported to reduce miRNA U-tailing (56), our study allows for the comparison of the relative contributions of all putative TENTs, demonstrating that other TENTs are not required. In particular, knockdown of *pup*-*2*, a paralog of *cid*-*1*, does not affect miRNA terminal uridylation (Figure S5).

Both CID-1 (also known as PUP-1 or CDE-1) and PUP-2 are orthologs of TUT4 and TUT7, which act redundantly, along with TENT2/TUT2/GLD2, to uridylate miRNA precursors in mammals (12, 15, 57). The common ancestor of nematodes and vertebrates likely contains only one TUT4/7 family gene, and a recent duplication occurred in each lineage to give rise to two paralogs (Figure 2C) (10). The duplication in nematodes likely occurred in the common ancestor of Rhabditina or earlier (Figure S7). In contrast, the *Drosophila* lineage has lost this gene family (10). Overall, the role of CID-1 demonstrates that the function of the TUT4/7 gene family in uridylation of miRNA species is deeply conserved, though the substrate specificity shows a different bias (preferentially targeting mature miRNAs rather than precursors in *C. elegans*) (Figure 2C).

To evaluate any redundancy between CID-1 and PUP-2, we assessed the U-tailing of miRNAs in individual knock-out mutants of *cid*-*1* and *pup*-*2* and a double *cid*-*1/pup-2* mutant. Three biological replicates of wild type and the single and double mutants were profiled by deep sequencing. The data from single mutant animals recapitulated the results from our RNAi screen. U-tailing was strongly disrupted in the *cid*-*1(tm1021)* mutant but not in the *pup*-*2(tm4344)* mutant (Figure 2D, Table S7). In the *cid*-*1/pup*-*2(om129)* double mutant, we observe a decrease in U-tailing that in most cases is comparable to that of the single *cid*-*1(tm1021)* mutant (Figure 2D, Table S7). This reaffirms the role of *cid*-*1* in U-tailing and suggests very little redundancy in its function with *pup*-*2*. Although CID-1 and PUP-2 are redundant in their biological function of protecting germline fate (58), they do not appear to be redundant for this molecular function. CID-1 and PUP-2 are much less similar (17.4%) than human TUT4 and TUT7 (44.5%), consistent with the lack of redundancy of CID-1 and PUP-2 for miRNA uridylation in *C. elegans.*

### miRNA adenylation requires F31C3.2/GLDR-2

We also analyzed the RNAi screen of putative TENTs to identify enzymes required for adenylated miRNA species, with the goal of assessing whether adenylation affects processing or stability of miRNAs. Because of the generally low level of A tailing and the noise inherent in tailing levels below 1%, we highlight those miRNAs that were >1% adenylated in empty vector samples in Figure 3. Knockdown of F31C3.2 reduced A-tailing of these miRNAs (*p*-value = 0.0003, Figure 3A-B, Figure S8) in multiple replicates (Figure S6C).

F31C3.2 is in the TENT2/GLD2 gene family of non-canonical cytoplasmic poly(A) polymerases, and we therefore name it <u>GLD</u>-2-<u>r</u>elated <u>2</u> (GLDR-2) (Figure 3C). The common ancestor of *Bilateria* likely had a single GLD2 gene, and duplications have occurred in the *Nematoda* lineage (GLD-2 and F31C3.2/ GLDR-2) and the *Drosophila* lineage (GLD2 and WISPY) (10). The nematode duplication likely occurred in the common ancestor of clades III-V or earlier (Figure S9). The single GLD2 gene in mammals and the paralog WISPY in *Drosophila* both catalyze adenylation of mature miRNAs (18, 19, 21, 47).

To determine whether F31C3.2/GLDR-2 has any functional redundancy with its paralog GLD-2, we assessed A-tailing in conditions where both genes are disrupted. We generated a F31C3.*2/gldr*-*2* knockout mutant, F31C3.2/*gldr*-*2*(*cdb187*), using CRISPR-Cas9, deleting the entire ORF of the gene. The mutant progeny were viable and superficially wild type. Because disruption of *gld-2* disrupts germline development and generates infertile animals (59), we used RNAi to knock down *gld-2* in a wild type or F31C3.2/*gldr*-*2(cdb187)* mutant background. Four biological replicates of each genotype/RNAi condition were deep sequenced. Results of single gene disruptions recapitulated those of the RNAi screen. A-tailing was significantly disrupted in the F31C3.2/*gldr*-*2(cdb187)* mutant on empty RNAi vector, but not in *gld*-*2* RNAi of wild type worms (Figure 3D, Figure S6D, Table S8). When *gld*-*2* RNAi was performed in the F31C3.2/*gldr*-*2(cdb187)* background, the disruption of A-tailing was comparable to that in the F31C3.2/*gldr*-*2(cdb187)* mutant on empty RNAi vector (Figure 3D, Figure S6D, Table S8). This suggests that there is little to no redundancy between F31C3.2/GLDR-2 and GLD-2 in miRNA tailing in *C. elegans*.

Here we show that the paralog F31C3.2/GLDR-2 - rather than GLD-2 itself - adenylates miRNAs in *C. elegans*, though the enrichment of this modification on 3p-derived miRNAs (Figure 1D) suggests that precursors rather than mature miRNAs are the substrate of F31C3.2/GLDR-2. Although F31C3.2/GLDR-2 previously had no known function *in vivo*, a high throughput assay in yeast showed that the enzyme is indiscriminate with respect to substrate nucleotide (60). However, knockdown of F31C3.2/GLDR-2 only affects the prevalence of A tailing (and not C, G, or U additions) in our *in vivo* experiments. Thus, additional factors in the *C. elegans* cellular context may confer nucleotide substrate specificity to F31C3.2/GLDR-2.

### RNAi screen does not identify nucleases targeting tailed miRNAs

In many previous studies, tailed miRNA precursors or tailed mature miRNAs are targeted to specific nucleases (61–64). To determine whether tailed miRNAs are the substrate of a specific nuclease, we also performed RNAi against a panel of nucleases previously implicated in miRNA turnover (4, 61, 62, 65–68). We observed only subtle effects on tailing (Figure 2A, 3A, S5 and S8). RNAi against exosome component *exos*-*4.1* showed statistically significant differences in U-tailing compared to the vector control (*p*-value = 0.016). However, the effect was a slight decrease in tailed species, suggesting an indirect effect of the RNAi. (The direct effect of knocking down an exonuclease that targets tailed miRNAs would be an increase in the prevalence of tailing.) Overall, these data suggest that tailed miRNAs may not be specifically targeted to any of the examined nucleases. (Note that PARN-1 was not included since previous studies showed no effect of *parn*-*1(lf)* on miRNA tailing or stability in *C. elegans* (69).) The knockdowns of nucleases displayed both a robust small RNA response mapping to each targeted locus and the expected deleterious phenotypes based on previous studies (70–73). Together this suggests that knockdown was sufficiently potent to impair activity of the enzymes. Since the strong pleiotropic phenotypes induced by these knockdowns introduce concerns of sample synchrony and staging, we did not analyze the impacts of these knockdowns on miRNA abundance. Overall, we have not observed evidence for the role of any of these nucleases in the trimming of tailed miRNAs.

### miRNAs are relatively long-lived

Having identified the enzymes required for miRNA tailing in *C. elegans*, we sought to examine the potential involvement of tailing in regulating miRNA turnover. To this end, we first characterized miRNA turnover in *C. elegans* by measuring miRNA half-lives using a system in which miRNA biogenesis can be acutely inactivated: a temperature-sensitive allele of the DGCR8 homolog *pash*-*1* (74). Upon upshift to restrictive temperature, *pash*-*1(ts)* no longer processes new miRNAs, and the decay of existing miRNAs can be observed over time (Figure 4A) (74, 75). To measure miRNA decay, we collected time course samples 0, 6, 24, and 48 hours after upshift to restrictive temperature. Because miRNAs comprise a large portion of the small RNA complement and global miRNA levels will change over the time course after *pash*-*1* inactivation, synthetic miRNAs not present in the *C. elegans* genome were spiked in to the samples at a constant concentration relative to total RNA to allow for robust normalization. Adult animals shifted to restrictive temperature produced a few non-viable progeny and did not proliferate further, so dilution of miRNAs by increasing amounts of total RNA is minimal in this system. After normalization to spike-ins, global decay of miRNAs was observed over the 48h time course in two biological replicates (Figure 4B, Table S9).

**Figure 4.**
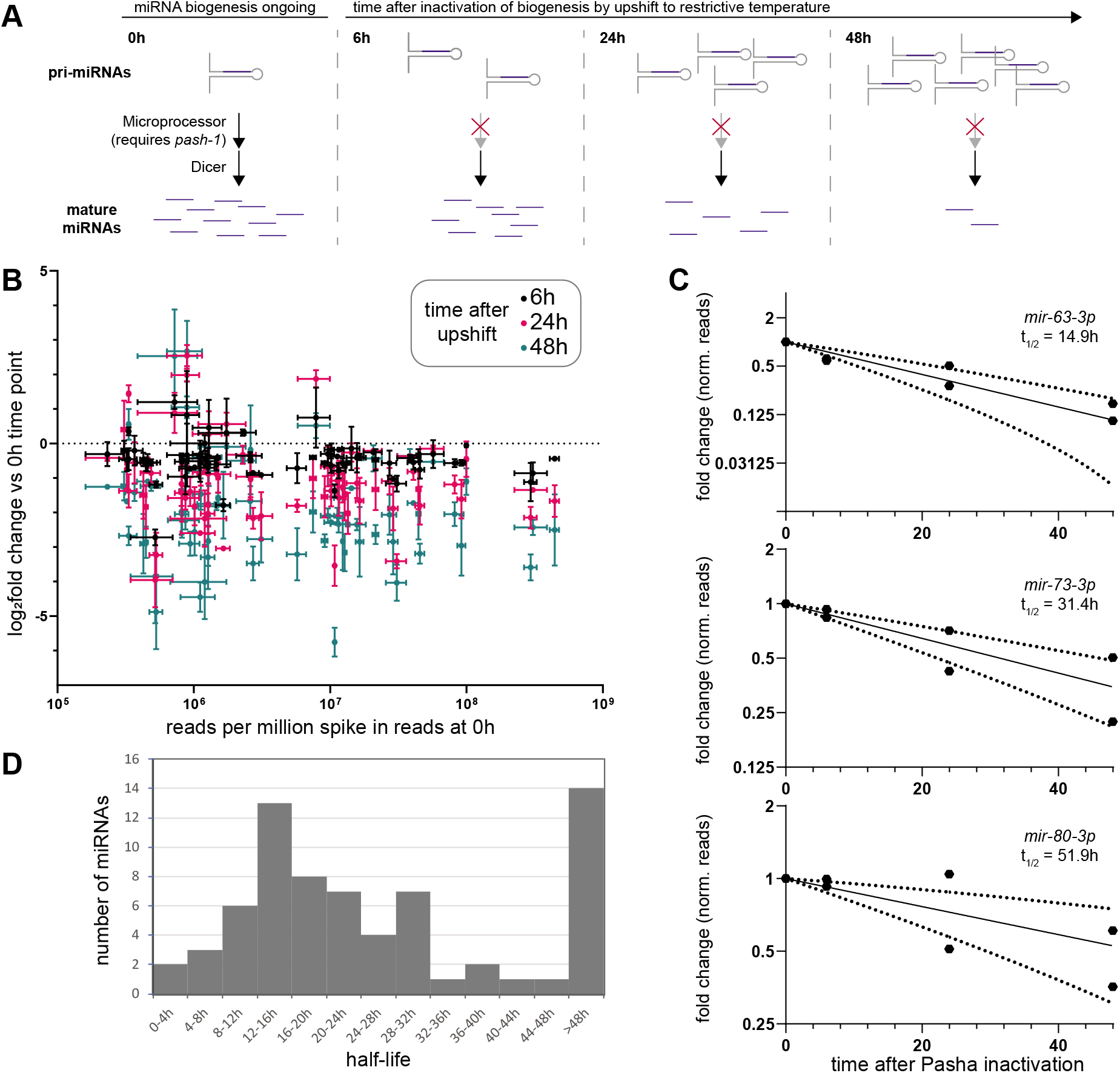
Characterization of miRNA half-lives in *C. elegans*. (A) Schematic of time courses after *pash*-*1* inactivation used for measuring miRNA decay. (B) Log_2_ fold change of miRNA reads compared to 0h after PASH-1 inactivation (y-axis) versus abundance at 0h (x-axis). Reads were normalized to spike-ins. Mean and standard error of two biological replicates are shown. (C) Representative time course data plotted as fold change, with exponential decay model and 95% prediction bands (dashed lines). Data for two biological replicates is shown. (D) Distribution of miRNA half-lives modeled from *pash*-*1(ts)* time course data. (B-D) Only miRNAs with >50 RPM in both replicates at the 0h timepoint were included in the analysis.

Using spike-in-normalized reads, time course data were fit to exponential decay curves to calculate miRNA half-lives. Half-lives determined using data from only one replicate were well correlated between replicates (Figure S10A). Therefore, we used data from both replicates together to fit half-life values with greater confidence. The fit of the decay curves was best when deep sequencing data were normalized to spike-ins, as expected, significantly outperforming normalization to total mapped reads (Figure S10B). Data for miRNAs with half-lives longer than the 48h course of the experiment (14 out of 68 analyzed miRNAs) generally did not fit the exponential decay curve well (Figure S10C, Table S10). In contrast, shorter-lived miRNAs fit the curve well, with 40 out of 54 miRNAs having a fit with R^2^> 0.8 (Figure 4C, Figure S10C, Table S10). Overall, miRNAs were fairly stable, with a median half-life of 20.7h (Figure 4D, Table S10), similar to most observations in mammalian and *Drosophila* cells (4, 6–8, 76). These results using spike-in normalization and deep sequencing yield longer half-lives than a previous study using *pash*-*1(ts)* (Figure S10D) (74). The previous study employed microarray miRNA quantification and normalization to non-miRNA small RNAs (siRNA and mirtron); when our data were normalized to mirtron reads, we observed a similar range of half-lives to their study (Figure S10E). This suggests that mirtron or siRNA normalization does not effectively control for library-wide changes. Nonetheless, both studies highlighted similar sets of fast-decaying miRNAs, with *mir*-*61*, *mir*-*71*, *mir*-*253*, and *mir*-*250* among the six lowest half-lives in both datasets (Table S10) (74).

### The proportion of tailed and trimmed miRNAs increases over time after biogenesis

Previous studies have demonstrated that terminal modifications correlate with miRNA turnover (22, 23, 26, 30, 31). These studies demonstrated that miRNAs that are targeted for decay display high levels of 3’ tailing. In our datasets, as miRNAs turn over during the time course after PASH-1 inactivation, the ensemble of miRNAs becomes increasingly skewed towards older miRNAs (approaching turnover). To determine whether miRNA terminal modifications are associated (correlatively or causatively) with miRNA turnover in our dataset, we quantified tailing across the time course. We observed that indeed later time points displayed higher levels of uridylation (Figure 5A-B, Table S9). This global trend included a large increase in tailing for *mir*-*70*-*3p* and many other miRNAs, although not all miRNAs displayed increased tailing over the time course (Figure 5C, Table S9). The increase of U-tailing on aging miRNAs is consistent with the mature miRNA being the substrate of U-tailing as suggested above (Figure 1C-D). No change was observed in adenylation of miRNAs over the time course (Figure 5A), consistent with its substrate being miRNA precursors.

**Figure 5.**
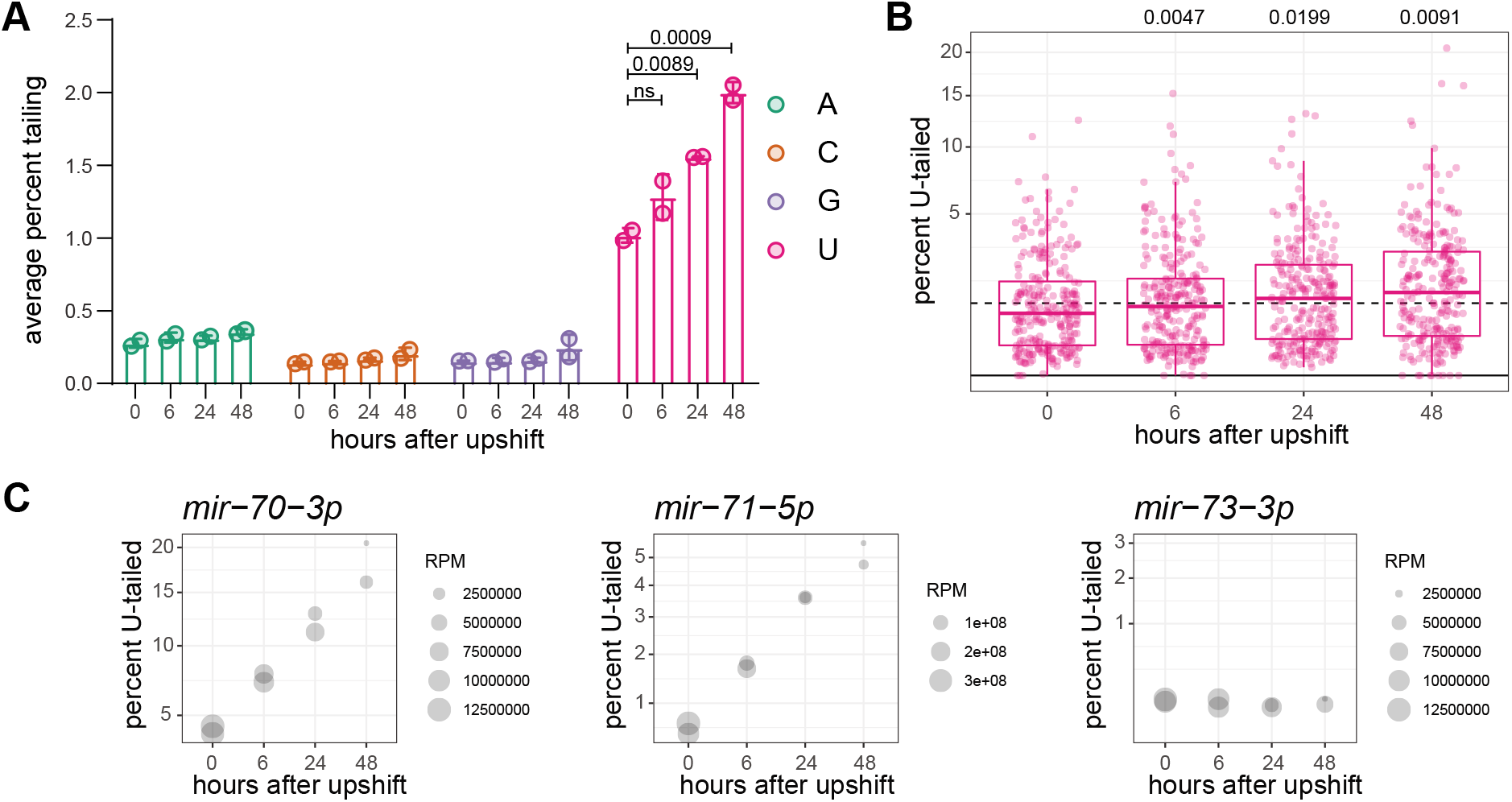
The proportion of tailed and trimmed miRNAs increases over time after biogenesis. (A) Average percent of reads bearing each indicated single-nucleotide tail. (B) Prevalence of mono-uridylation of miRNAs across time course after PASH-1 inactivation in one biological replicate. Each dot represents an individual miRNA. (A-B) One-way ANOVA was used to compare all time points to 0h; when ANOVA was significant, Dunnett’s multiple comparison test was performed, and *p*-values are shown above graph. (C) Representative plots for indicated miRNAs, where y-axis indicates percent mono-uridylation, and size of dot represents read number. Note that reads here are reads per million spike-in reads, an arbitrary unit.

Like tailing, 3’ trimming is also often observed concomitant with tailing (22, 26). Therefore, we also analyzed trimming over the time course after PASH-1 inactivation. Like tailing, global 3’trimming increased in later time points, as indicated by a lower proportion of canonical-length reads and increased proportion of shorter genome-matching reads (Figure S11). Like tailing, this subtle global trend was more apparent when examining individual miRNAs (Figure S11). Thus, both tailing and trimming increase in older miRNAs that are approaching turnover.

If miRNA tailing regulates turnover, two models may explain the increase in U-tailing observed over the time course after PASH-1 inactivation. On one hand, if tailing triggers turnover, then older miRNAs would be tailed as they approach decay, increasing the relative abundance of tailed species over time. On the other hand, if tailing protects miRNAs from decay, this could also result in the relative accumulation of tailed miRNAs at later time points, as non-tailed miRNAs are decayed more rapidly.

We therefore asked whether tailing is correlated (or anti-correlated) with the rate of miRNA turnover. We did not observe an overall correlation between initial prevalence of the U-tailed isoform and half-life for a given miRNA (Figure S12A), nor between rate of increase in tailing and half-life (Figure S12B). While this indicates that terminal uridylation is unlikely to be the primary determinant of turnover, uridylation might play a modulatory role that is not apparent in bulk analysis due to larger effects of other unknown parameters.

### Abrogation of miRNA tailing does not affect miRNA turnover

To determine whether uridylation may play a modulatory role - either stimulating or impairing turnover - we sought to disrupt miRNA tailing and then measure rates of miRNA turnover in its absence. In wild type animals, disrupting U-tailing in the context of *cid*-*1* RNAi, *cid*-*1* knockout, or *cid*-*1/pup*-*2* double knockout did not significantly change abundance of miRNAs, suggesting that turnover is unchanged (Figure 6A-C). To further examine the effect of tailing on miRNA decay, we profiled in *cid*-*1* RNAi after PASH-1 inactivation in the *pash*-*1(ts)* background (0 and 24h after upshift). RNAi against *cid*-*1* reduced U-tailing at both time points (Figure S13A, Table S11). However, when we examined the change in miRNA abundance from zero to 24h after PASH-1 inactivation, *cid*-*1* RNAi did not have a global effect. We divided the miRNAs into subsets based on the percent of tailed species and saw no effect on turnover in miRNAs that were tailed 0-1%, 1-2%, or 2-30% (Figure 6D). One exception to this was miRNAs of the embryo-enriched *mir*-*35*-*41* cluster, which showed a higher log_2_fold change in *cid*-*1* RNAi than vector control (Figure 6E). However, this is likely due to a lower starting amount of these miRNAs in T0 *cid*-*1* RNAi samples (Figure 6F); indeed, these miRNAs are the only ones with less than half the abundance in *cid*-*1* RNAi T0 samples as compared to vector T0 samples. Furthermore, both strands of the miRNA duplex derived from these precursors behaved similarly, supporting an effect of *cid*-*1* RNAi on transcription or biogenesis rather than on decay rate of the mature miRNA species (Figure 6E-F). Because these miRNAs are largely germline and embryo-specific (53, 77), the difference in their abundance most likely corresponds to slight differences in staging of *cid*-*1* RNAi T0 samples due to pleiotropic effects of *cid*-*1(lf)*, which negatively impacts germline function and embryo production (58). Finally, we also specifically examined miRNAs that show high increases in uridylation after *pash*-*1* inactivation to determine whether these cases might be particularly sensitive to disrupting uridylation by *cid*-*1* RNAi. For these miRNAs, tailing was disrupted, but turnover was not reproducibly altered (Figure 6G). Taken together, these data show that miRNA uridylation in *C. elegans* is not a global regulator of miRNA decay.

**Figure 6.**
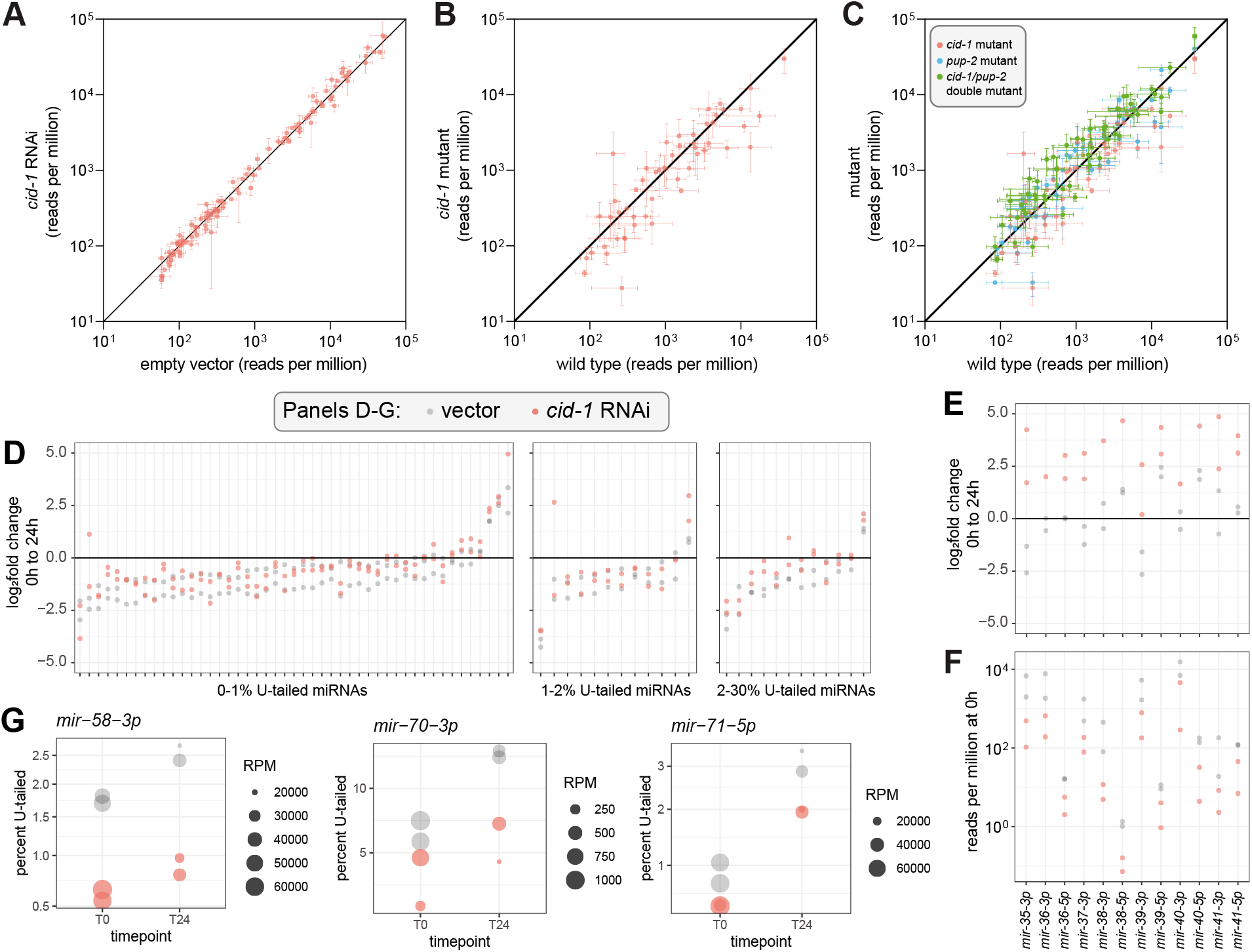
Uridylation does not globally regulate miRNA abundance or decay. (A) Abundance of miRNAs in *cid*-*1* RNAi versus empty vector. Mean and standard error are plotted. (B-C) Abundance of miRNAs in indicated mutant versus wild type. (A-C) Except for *cid*-*1* RNAi (two biological replicates), mean and standard error of three biological replicates are plotted. All miRNAs >50 RPM in all empty vector or wild type replicates, respectively, are shown. (D) Log_2_fold change in reads per million from zero to 24h after PASH-1 inactivation. miRNAs are divided into three plots by the average percent mono-uridylation in empty vector at 0h. All miRNAs with >50 RPM at 0h in both empty vector replicates is shown, except for *mir*-*35*-*41*. (E) Log_2_fold change from zero to 24h after PASH-1 inactivation for *mir*-*35*-*41* guide (3p) strands and star (5p) strands. All miRNAs from this group are shown, regardless of abundance, except for *mir*-*35*-*5p* and *mir*-*37*-*5p* which had ~0 reads in all samples. (F) Abundance of *mir*-*35*-*41* guide and star strands at 0h (prior to PASH-1 inactivation). (G) Plots showing percent mono-uridylation and miRNA abundance at zero and 24h after PASH-1 inactivation. Size of dot represents abundance, and y-axis is percent mono-uridylation.

Next, we examined whether A-tailing altered miRNA abundance or turnover. In wild type, although F31C3.2/*gldr*-*2* RNAi reduces A-tailing, RNAi did not alter miRNA abundance (Figure S14A-B). F31C3.2/*gldr*-*2* knockout and F31C3.2/*gldr*-*2* knockout with *gld*-*2* RNAi also showed minimal effects on miRNA abundance (Figures S14C-F). Small changes in the abundance of three *mir*-*35*-*42* family members appeared to be due to changes in synthesis, rather than changes in turnover of the miRNAs, as indicated by similar changes in star strand abundance (Figure S14G); as noted above, this is likely due to slight staging differences due to pleiotropic effects. In the *pash*-*1(ts)* setting, fewer miRNAs could be examined with high confidence due to lower abundance and/or lower A-tailing. (Lower miRNA abundance may be the result of reduction of *pash*-*1* function even at permissive temperature, and likewise, a reduced lifetime of miRNA precursors in this setting could lead to reduced A-tailing.) Nonetheless, we still observed that F31C3.2/*gldr*-*2* RNAi abrogated A-tailing in the 24h time point (Figure S14H, Table S11). When examining the change in miRNA abundance from zero to 24h after temperature shift, F31C3.2/*gldr*-*2* RNAi did not affect the rate of miRNA decay (Figure S14I). Therefore, the adenylation mediated by F31C3.2/GLDR-2 does not regulate miRNA stability.

### High levels of mirtron uridylation do not affect abundance

Mirtrons are non-canonical miRNAs that derive from small introns; debranching of these intron lariats results in a miRNA precursor-like hairpin structure, bypassing the requirement for Microprocessor-mediated cleavage (78, 79). Previous work showed that mirtron precursors are highly uridylated and that, in *Drosophila*, this uridylation is carried out by the *Drosophilidae*-specific TUTase6-related enzyme Tailor and negatively regulates mirtron abundance (10, 16, 17, 80). In wild type sequencing libraries, only one mirtron, *mir*-*62*, was abundant enough for reliable tailing analysis. However, in *pash*-*1(ts)* libraries at restrictive temperature, mirtrons are relatively elevated in sequencing libraries since they do not depend on Microprocessor activity, while canonical miRNAs are globally depleted. This allowed for the analysis of tailing of multiple mirtrons. We observed that (as previously reported from aggregate sequencing data), 3p-derived mirtrons are highly uridylated to a much greater degree than canonical miRNAs, with the exception of *mir*-*62* (Figure S13B) (80). Not only is *mir*-*62* atypical in its proportion of tailing, but *mir*-*62* is also much more abundant than the other mirtrons (Figure S13B). Like uridylation of canonical miRNAs, uridylation of mirtrons also required CID-1 (Figure S13B). Surprisingly, although knockdown of *cid*-*1* strongly abrogated mirtron tailing, the abundance of the mirtrons was not elevated as was observed in *Drosophila* (upon knockdown of Tailor) (Figure S13B). Therefore, although the high rate of U-tailing is conserved for most *C. elegans* mirtrons, this modification does not regulate mirtron abundance.

### Standard over-expression transgenes unlikely to induce TDMD in *C. elegans*

Recent work provided evidence that some *C. elegans* miRNAs are targeted by TDMD, whereas another previous study described <u>t</u>arget-<u>m</u>ediated <u>m</u>iRNA <u>p</u>rotection (TMMP) of miRNAs in *C. elegans* (35, 81). To further interrogate the relationship between target binding and miRNA stability in *C. elegans*, we attempted to induce TDMD of specific miRNAs using exogenous transgenic triggers. We modified a ubiquitously expressed GFP construct (*Peft*-*3::GFP::tbb*-*2* 3’UTR) to contain two highly complementary binding sites for a miRNA of interest in the transgene’s 3’UTR (Figure S15A). We chose to target *lin*-*4*, *mir*-*237*, and *mir*-*85* because of their moderate expression level, moderate to slow decay rate, and low proportion of tailing. Also, the biology of *lin*-*4* is well studied, and it shares a seed sequence with *mir*-*237* (82). Extra-chromosomal arrays expressing these transgenes did not induce any phenotype (such as a *lin*-*4* loss-of-function phenotype). Deep sequencing showed that the miRNAs were not altered in abundance in the presence of the transgene expressing their cognate putative TDMD trigger (Figure S15B). The TDMD triggers also did not alter tailing of the targeted miRNAs (Figure S15C). Most previously described triggers of TDMD are expressed at the same or higher levels than the targeted miRNA (23, 24, 29); the most efficient known TDMD trigger, the lncRNA Cyrano, is an extreme case which effects multiple turnover TDMD when expressed at a 17-fold lower level than its target miRNA (31). To determine if our transgenes are expressed at a similar stoichiometry with respect to the targeted miRNAs, we performed absolute quantification of the transgene mRNA and the targeted miRNAs using digital droplet PCR (ddPCR) and miRNA Taqman assays, respectively. Despite being expressed from a high-copy transgene with an efficient promoter, each transgene mRNA was >300-fold lower in abundance than the targeted miRNA (Figure S15D). Thus, the stoichiometry renders these transgenes unlikely to induce TDMD. We are so far unable to dissect the features of TDMD in *C. elegans* using traditional protein-coding transgenes as synthetic triggers.

## Discussion

In this work, we characterize the nature of miRNA tailing in *C. elegans*, with the broad view of determining ancestral functions of this class of post-transcriptional modification. We show that uridylation is far more prevalent than adenylation in *C. elegans*, unlike mammals and *Drosophila* in which both modifications are common (46–49).

We show that the substrate of these additions is shifted in different lineages. Based on its even distribution between 3p- and 5p-derived miRNAs, we infer that uridylation targets mature miRNAs in *C. elegans*, whereas precursors are the predominant substrate in mammals and *Drosophila* (33, 46–48, 51). In contrast, adenylation is skewed towards 3p-derived miRNAs in *C. elegans*, indicating that miRNA precursors are its likely substrate, unlike in vertebrates where mature miRNAs are more often adenylated and *Drosophila* where both precursors and mature miRNAs are likely substrates (46–48).

Despite the shifts in the apparent substrate of the modification, the enzymes required for these modifications are largely conserved. MiRNA uridylation requires CID-1, which is in the same TUT-4/7 gene family that carries out this modification (along with TUT2/GLD2) in mammals (12, 15, 57). MiRNA adenylation requires F31C3.2/GLDR-2. F31C3.2/GLDR-2 is in the TUT2/GLD2/PAPD-4 gene family which carries out miRNA adenylation in mammals and *Drosophila* (18, 19, 21, 47). Importantly, the enzymatic activities of these enzymes in uridylation/adenylation has been previously demonstrated *in vitro* or in a heterologous tethering assay (56, 60).

We characterize rates of miRNA decay in *C. elegans*, which are largely similar to those in other organisms (4, 6–8, 76). The median half-life – 20.7h – is relatively long. Half-lives in this study may even be slightly underestimated due to slight expansion of the culture after upshift to restrictive temperature. Nonetheless, certain miRNAs (especially *mir*-*61*, *mir*-*71*, *mir*-*253*, and *mir*-*250*) are reproducibly short-lived across replicates in our study and previous studies (74). How these fast decay rates are specified will be an important area of ongoing study. Moreover, we have only analyzed a single condition; decay may be differentially regulated for subsets of miRNAs in distinct developmental stages or under stress conditions. Notably, a study conducted in L1 stage larvae highlighted different fast-decaying miRNAs than our study in adults (83), suggesting that miRNA turnover may be developmentally regulated.

Although we observed that the proportion of tailed species increased over time after biogenesis, disrupting tailing had no impact on turnover. This is consistent with recent kinetic analysis which demonstrated that tailing is faster than 3’ trimming, so miRNAs are more likely to carry a tail as they approach decay (8). TDMD is the most frequently-studied context in which tailing correlates with decay, yet recent work on TDMD suggests that tailing is correlative – not causative – of TDMD, and that conformational changes that promote decay of the Argonaute-miRNA complex also expose the miRNA 3’ end to TENTs (31, 32, 34, 35). However, in specific developmental contexts, tailing does promote wholesale miRNA turnover (18). Therefore, cell-type or developmental contexts which we have not yet examined may also couple decay to tailing in *C. elegans*. By identifying the enzymes required for miRNA tailing, this study lays the groundwork for discovering and dissecting such regulation.

## Data Availability

All raw sequence data will be deposited in NCBI Sequence Read Archive (BioProject Number PRJNA703306). Reviewer link: https://dataview.ncbi.nlm.nih.gov/object/PRJNA703306?reviewer=2h12a5i72qildfkt3l259bdli9

## Acknowledgments

We thank WormBase, the NIDDK Genomics Core, the NCI CCR Genomics Core, and NIH High Performance Computing. Strains are regularly received from the CGC, which is funded by NIH Office of Research Infrastructure Programs (P40 OD010440). Thank you to Anna Zinovyeva and members of the McJunkin lab for helpful discussions and to Cameron Palmer for bioinformatics support.

