## Supplemental Figures S1-S15 for "Screening by deep sequencing reveals mediators of microRNA tailing in *C. elegans*"

**List of Supplemental Materials**

**Figure S1.** Correlation of miRNA tailing across biological replicates.

**Figure S2.** miRNA tailing across development.

**Figure S3.** Abundance of highly tailed miRNAs across development.

**Figure S4.** Tailing of highly tailed miRNA species across development.

**Figure S5.** Mono-uridylation of miRNAs in RNAi screen of candidate trimming and tailing enzymes.

**Figure S6.** Independent experiments confirm roles of *cid-1* and F31C3.2 in miRNA tailing.

**Figure S7.** Phylogenetic relationship of CID-1, PUP-2, TUT4, and TUT7.

**Figure S8.** Mono-adenylation of miRNAs in RNAi screen of candidate trimming and tailing enzymes.

**Figure S9.** Phylogenetic relationship of GLD-2 and F31C3.2.

**Figure S10.** Summary data from half-life calculations.

**Figure S11.** Trimming of miRNAs over time course after PASH-1 inactivation.

**Figure S12.** Lack of correlation of half-life with mono-uridylation.

**Figure S13.** Impact of untemplated uridylation on miRNA abundance and stability.

**Figure S14.** Impact of untemplated adenylation on miRNA abundance and stability.

**Figure S15.** Standard over-expression transgenes unlikely to induce TDMD in *C. elegans*.

**Table S1.** Sources of RNAi clones.

**Table S2.** Oligonucleotides and strains used in this study.

**Table S3.** Small RNA sequencing library sample IDs and number of total and genome-mapped reads.

**Table S4.** Percent single-nucleotide tails in wild type adults.

**Table S5.** Abundance and percent single-nucleotide tails in published developmental time courses.

**Table S6.** Abundance and tailing of miRNAs in RNAi screen.

**Table S7.** Abundance and tailing of miRNAs in *cid-1* and *pup-2* and *cid-1/pup-2* mutant strains.

**Table S8.** Abundance and tailing of miRNAs in wild type and F31C3.2/*gldr-2* mutant backgrounds, on empty vector or *gld-2* RNAi.

**Table S9.** Abundance and tailing of miRNAs across time course after PASH-1 inactivation.

**Table S10.** Results of one phase decay non-linear regression to estimate miRNA half-lives.

**Table S11.** Abundance and tailing of miRNAs in *pash-1(ts)* animals before and 24h after upshift to restrictive temperature.

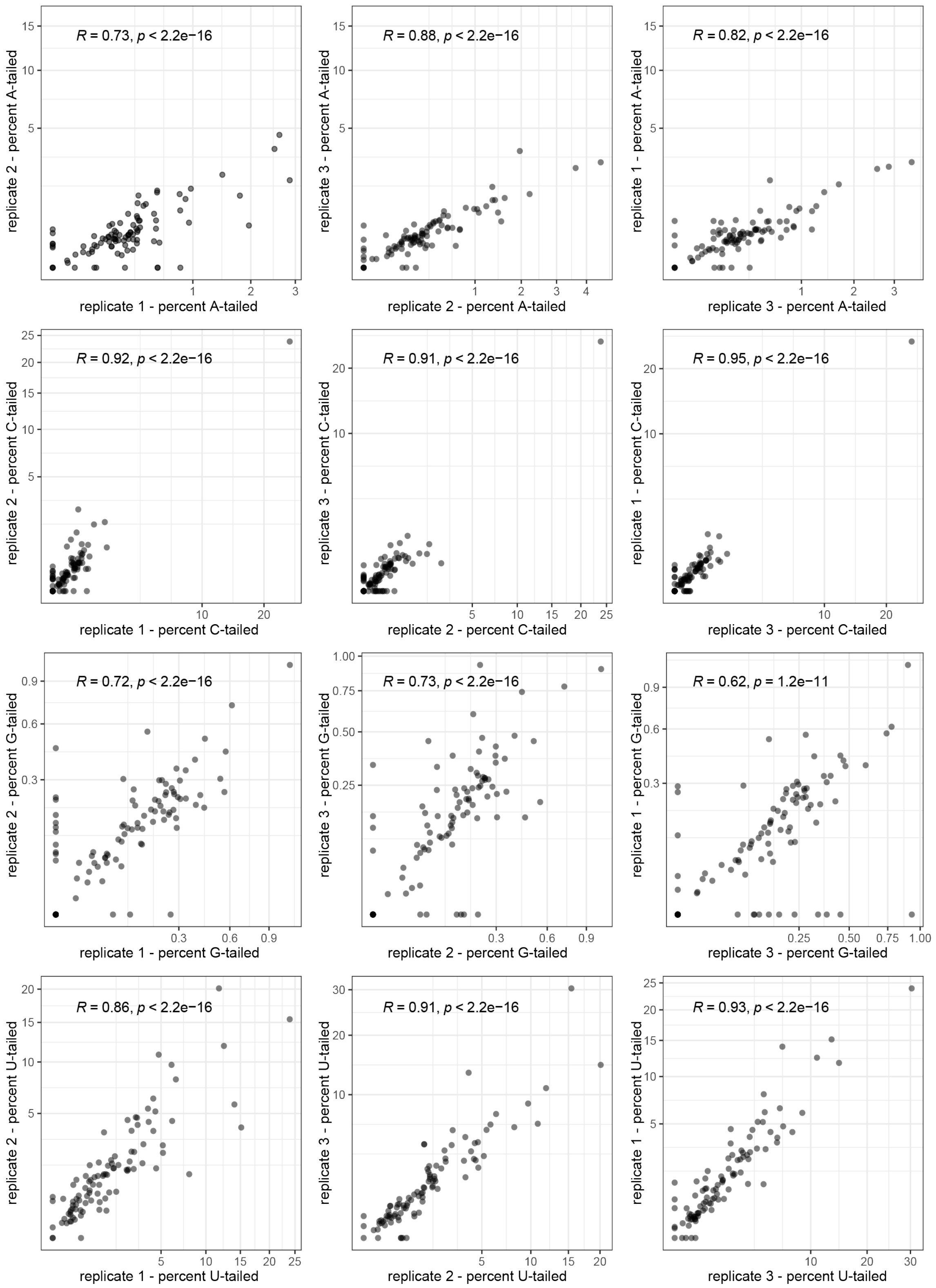


**Figure S1. Correlation of miRNA tailing across biological replicates.** Pearson's correlation test r and p-values are shown. Only miRNAs with > 50 RPM in all biological replicates are plotted.

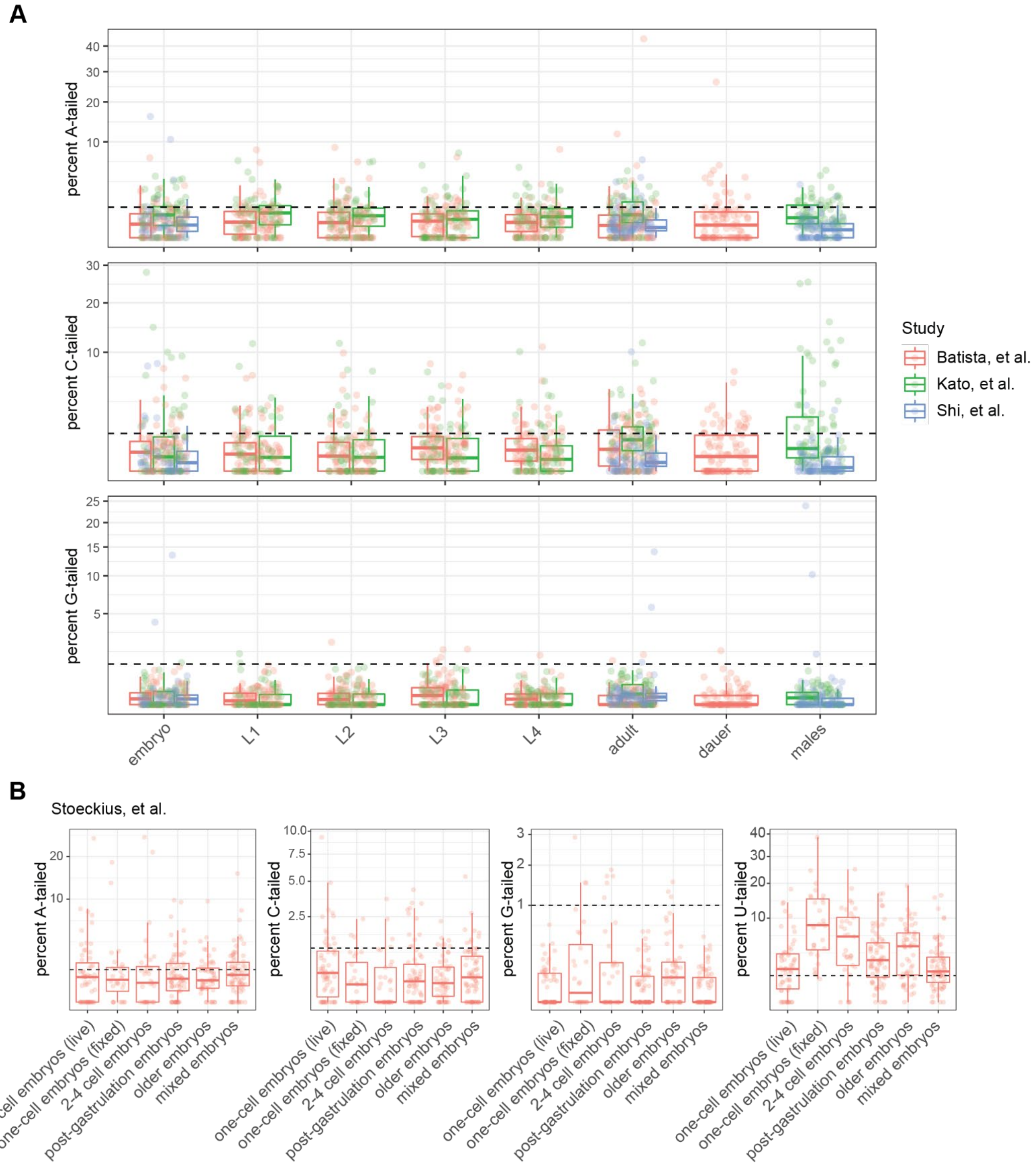


**Figure S2. miRNA tailing across development.** (A) Meta-analysis of three published datasets shows tailing across *C. elegans* developmental stages. Percent of each miRNA bearing the indicated single nucleotide tail is shown. No global changes in tailing are observed. See Figure 1E for uridylation. (B) miRNA tailing in staged embryos does not show consistent global regulation of tailing. (A-B) miRNAs are shown if they are >50 RPM in the indicated sample. See main text for full citations of analyzed datasets.

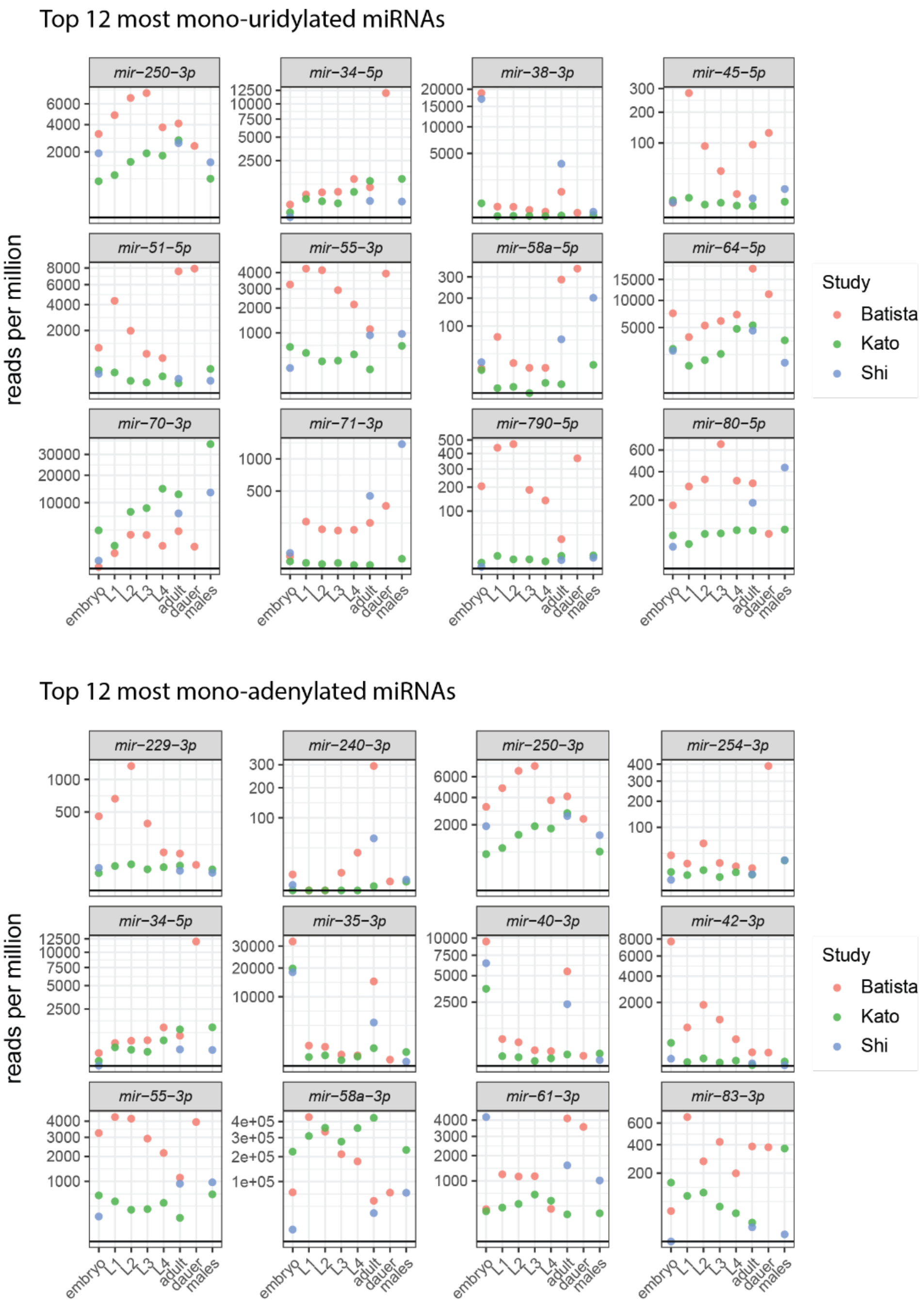


**Figure S3. Abundance of highly tailed miRNAs across development.** Abundance of miRNAs across development from three prior studies. Top twelve mono-uridylated or mono-adenylated miRNAs were selected based on mean of three biological replicates in early gravid adults from this study.

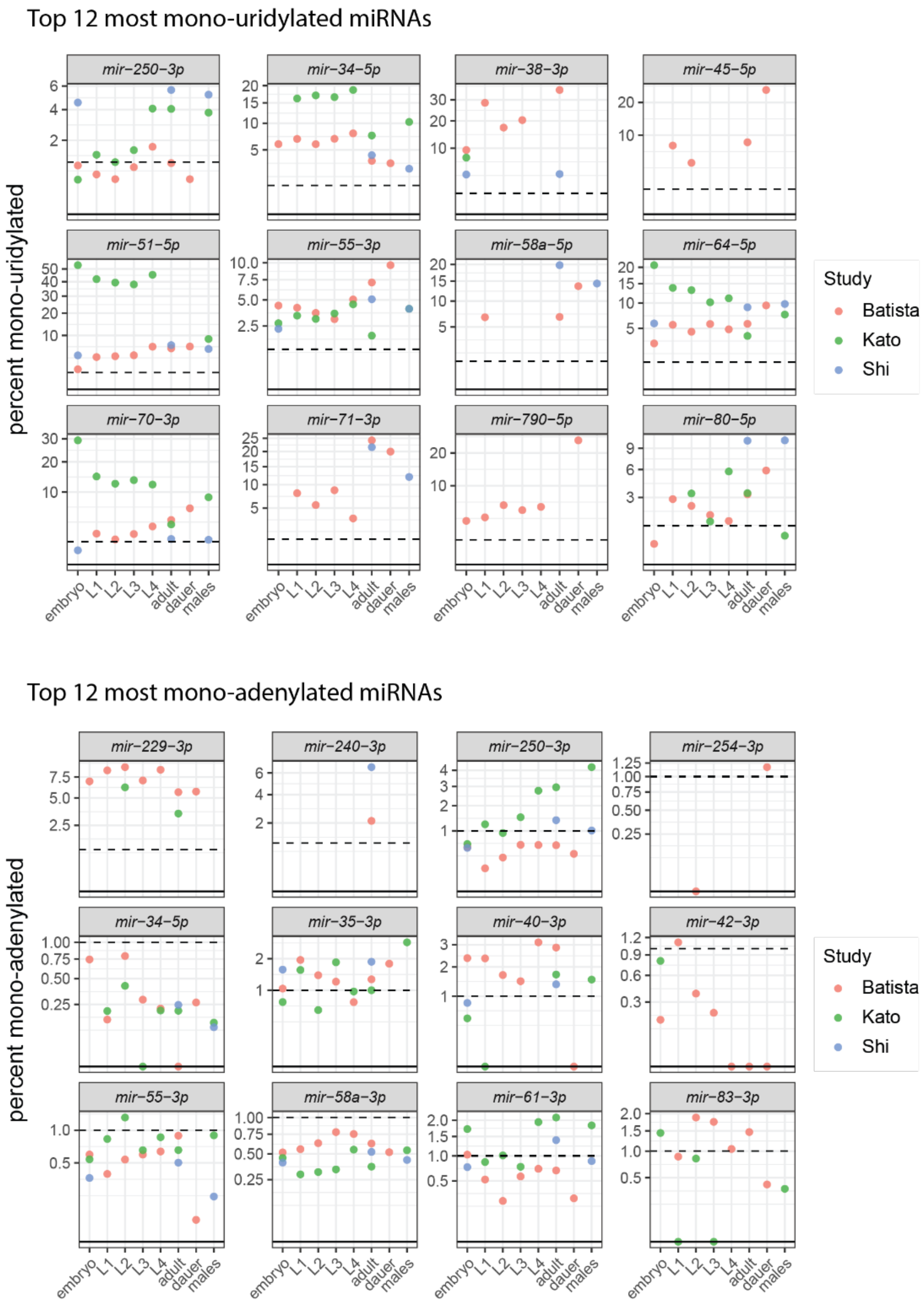


**Figure S4. Tailing of highly tailed miRNA species across development.** Percent of miRNA tailing across development from three prior studies. Top twelve mono-uridylated or mono-adenylated miRNAs were selected based on mean of three biological replicates in early gravid adults from this study. Data is only shown for samples in which the miRNA is >50 RPM.

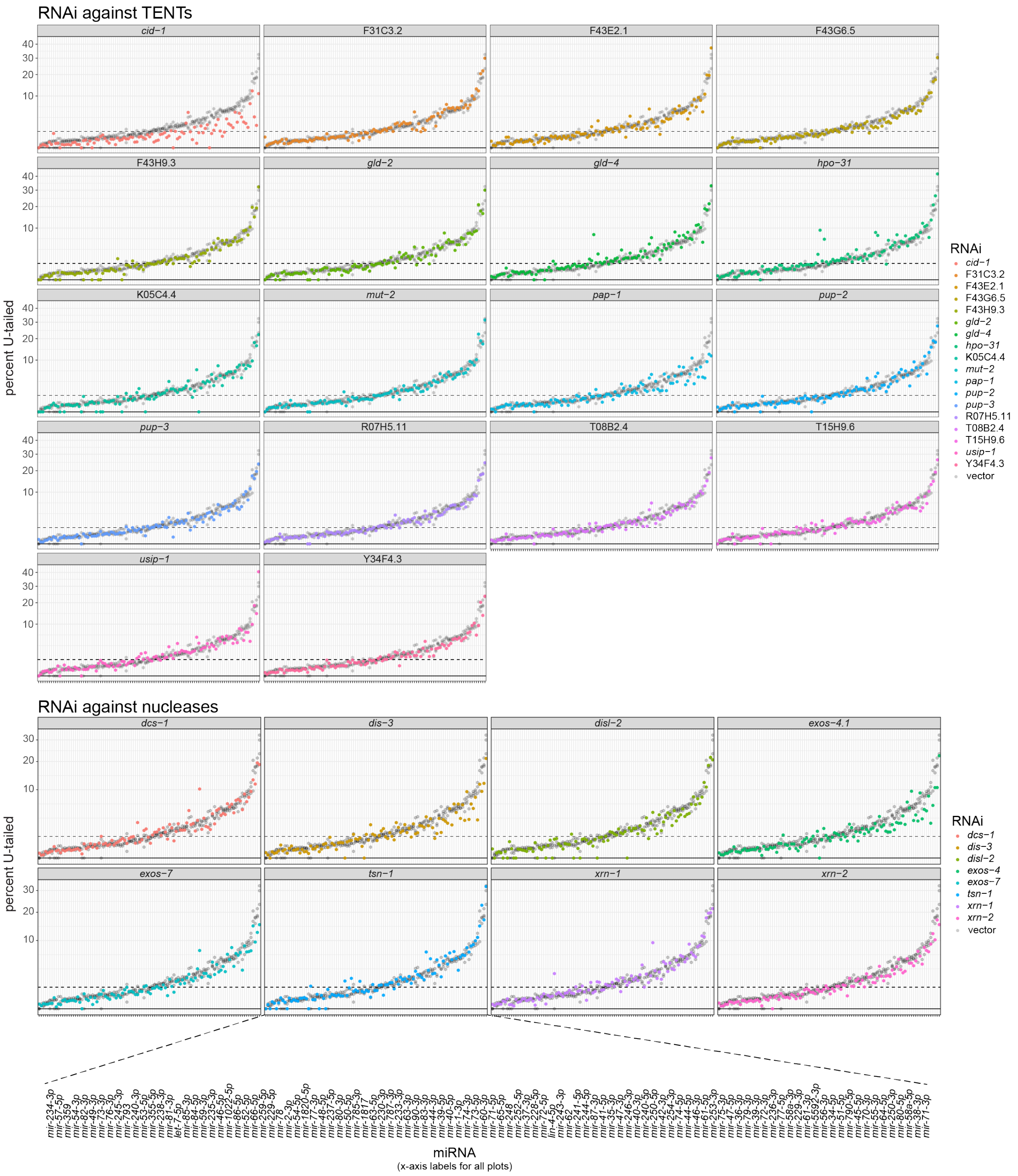


**Figure S5. Mono-uridylation of miRNAs in RNAi screen of candidate trimming and tailing enzymes.** Percent mono-uridylation in vector or indicated RNAi. Each column is an individual miRNA. All miRNAs with >50 RPM in all empty vector replicates are shown. Three biological replicates of empty vector are shown in gray on all plots. Colored dots correspond to RNAi condition indicated above graph. Top: RNAi against candidate tailing enzymes. Bottom: RNAi against candidate nucleases.

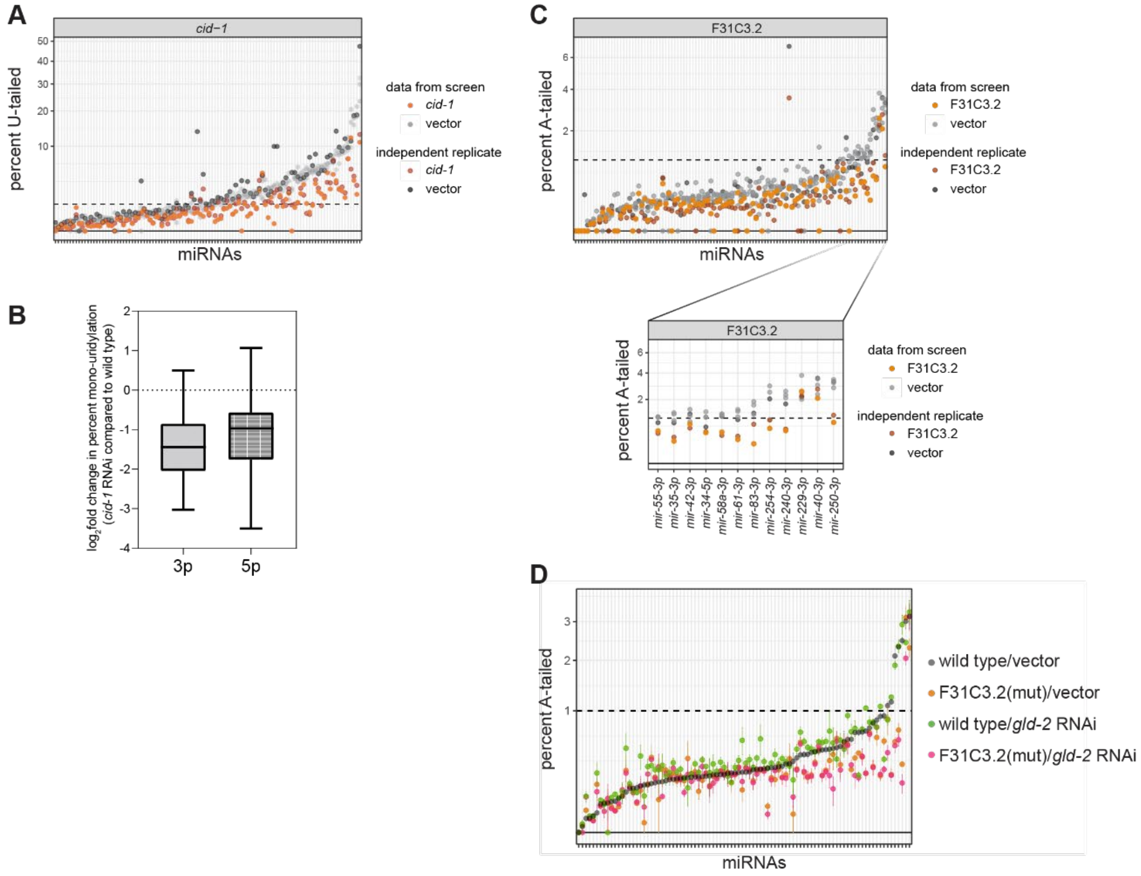


**Figure S6. Independent experiments confirm roles of *cid-1* and F31C3.2 in miRNA tailing.** (A) Percent mono-uridylation in vector or *cid-1* RNAi. Data from Figure 2B are re-plotted (data from screen) along with data from an independent biological replicate. (B) Box plot of $\log_2$fold change in percent mono-uridylation in *cid-1* RNAi compared to wild type, separated by 3p or 5p origin of the miRNA. No significant difference was observed between the two groups. (C) Percent mono-adenylation in vector or F31C3.2/*gldr-2* RNAi. Data from Figure 3B are re-plotted (data from screen) along with data from an independent biological replicate. (D) Percent mono-adenylation in vector or *gld-2* RNAi in a wild type or F31C3.2/*gldr-2* mutant background. Mean and standard error of four biological replicates is shown. (Same data as plotted in Figure 3D, but miRNAs with <1% tailing in wild type are included.) (A,C-D) Each column is an individual miRNA. All miRNAs with >50 RPM in all wild type empty vector replicates are shown.

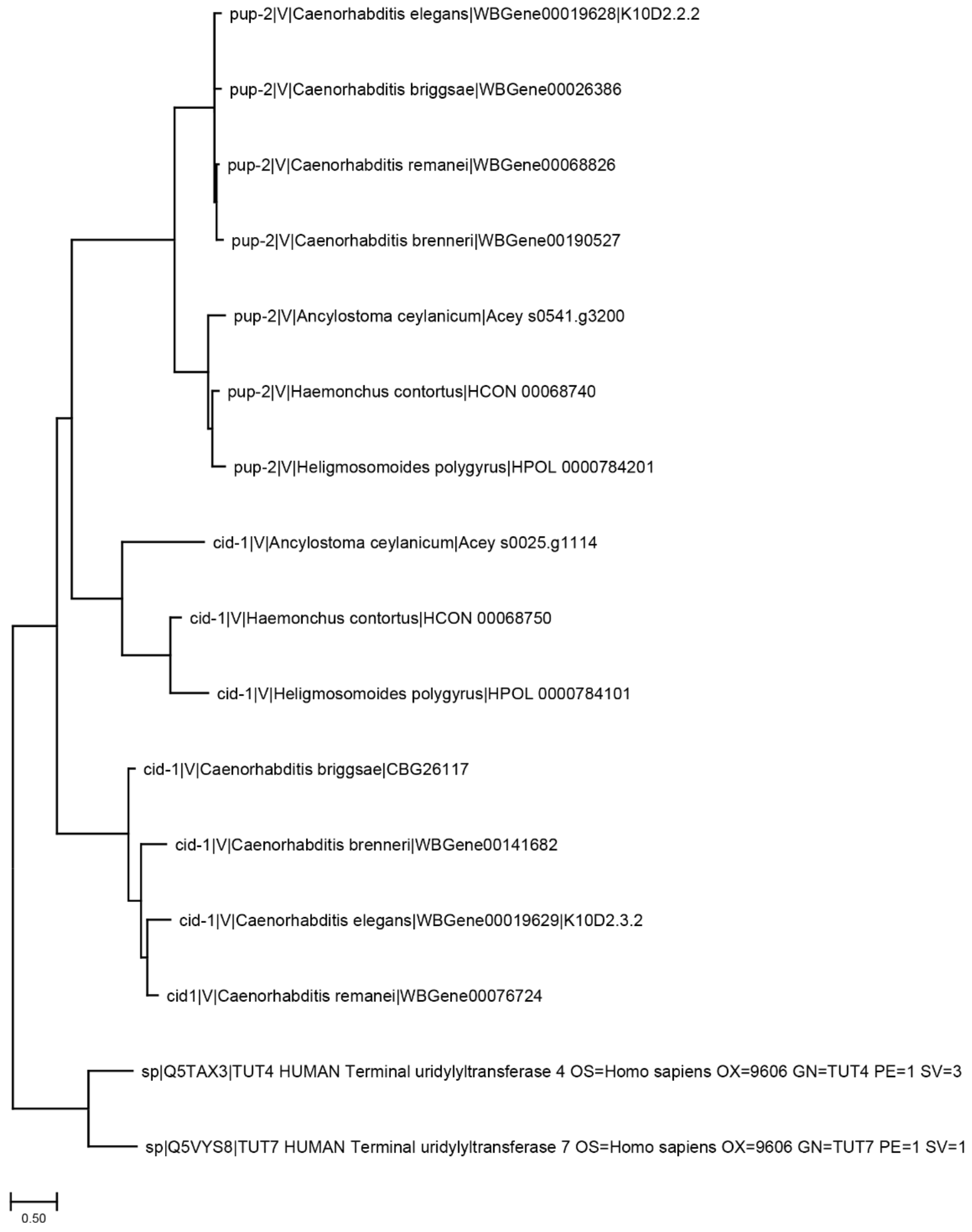


**Figure S7. Phylogenetic relationship of CID-1, PUP-2, TUT4, and TUT7.** The roman numeral following the gene name indicates the nematode clade containing the indicated species (WormBase ParaSite version WBPS15). Very few species outside of clade V had strong orthologs for *C. elegans* CID-1 and PUP-2. Inside clade V, hits were found for subclades Strongyloidea and Rhabditoidea; we included orthologues from three species and in each sub-group, as well as the human orthologs. No orthologs were identified in Diplogasteromorpha (Pristionchus).

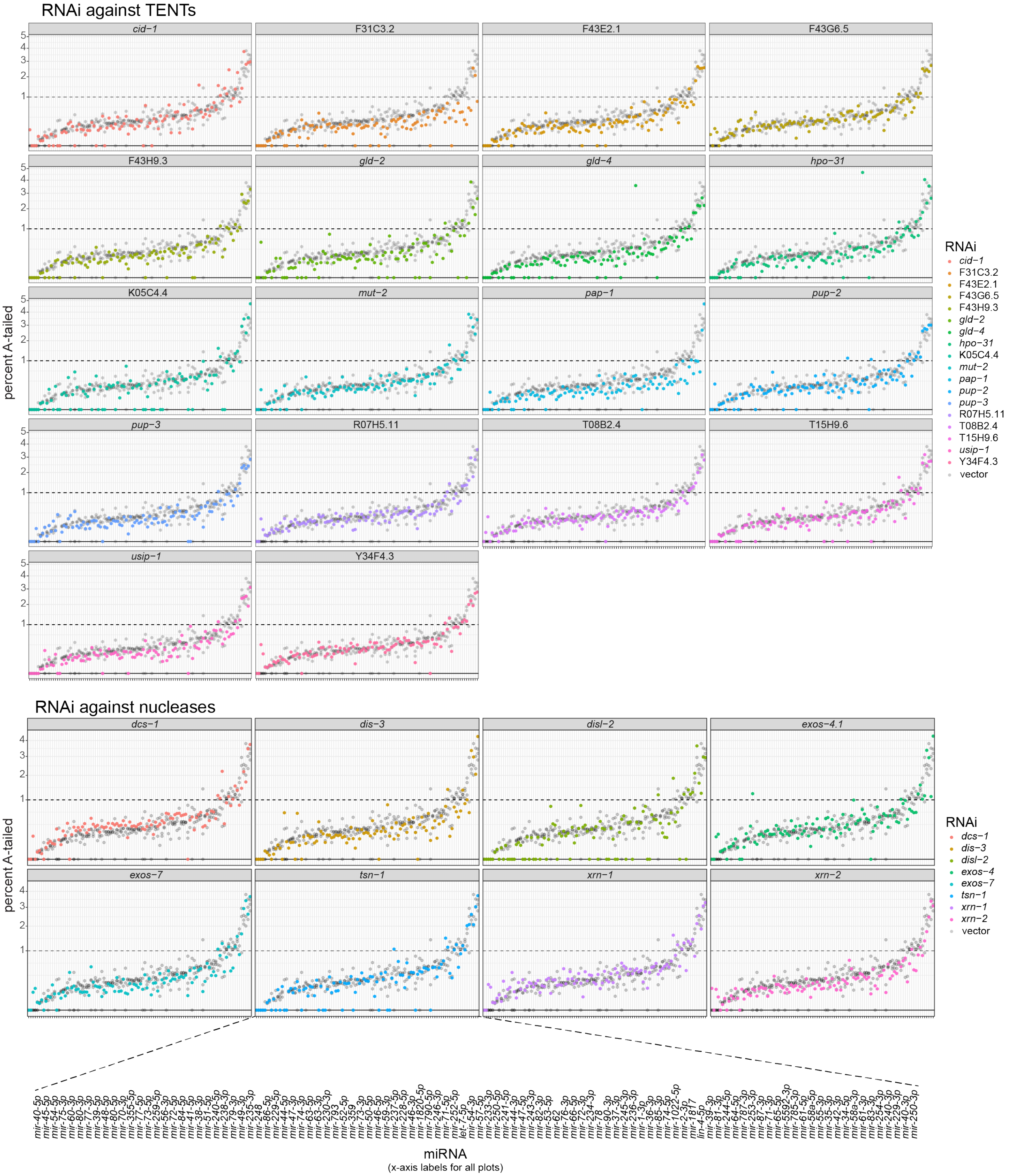


**Figure S8. Mono-adenylation of miRNAs in RNAi screen of candidate trimming and tailing enzymes.** Percent mono-adenylation in vector or indicated RNAi. Each column is an individual miRNA. All miRNAs with >50 RPM in all empty vector replicates are shown. Three biological replicates of empty vector are shown in gray on all plots. Colored dots correspond to RNAi condition indicated above graph. Top: RNAi against candidate tailing enzymes. Bottom: RNAi against candidate nucleases.

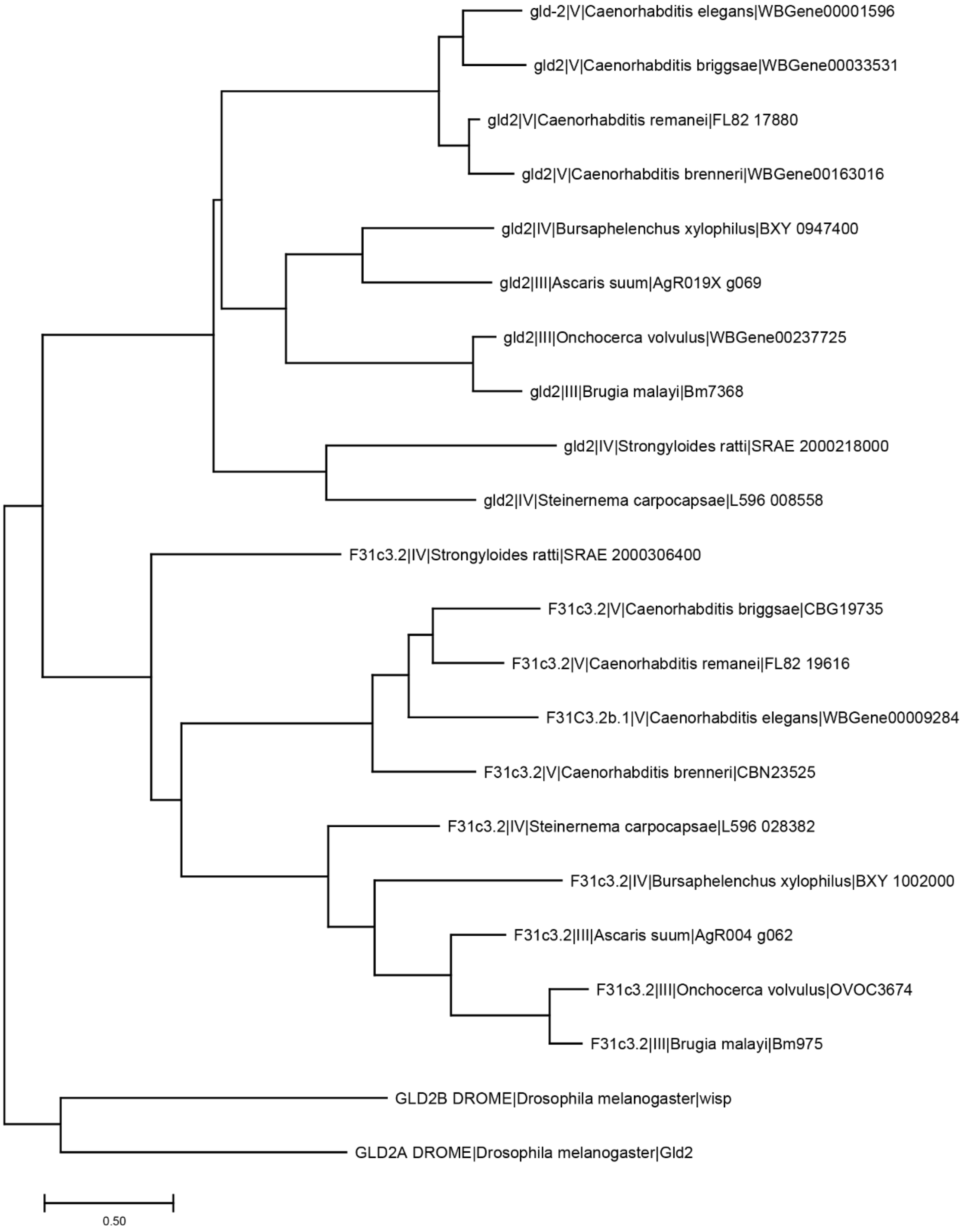


**Figure S9. Phylogenetic relationship of GLD-2 and F31C3.2.** The roman numeral following the gene name indicates the nematode clade containing the indicated species (WormBase ParaSite version WBPS15). In addition to *C. elegans* sequences, orthologs were included for three species from each of clades III, IV, and V, as well as *D. melanogaster*.

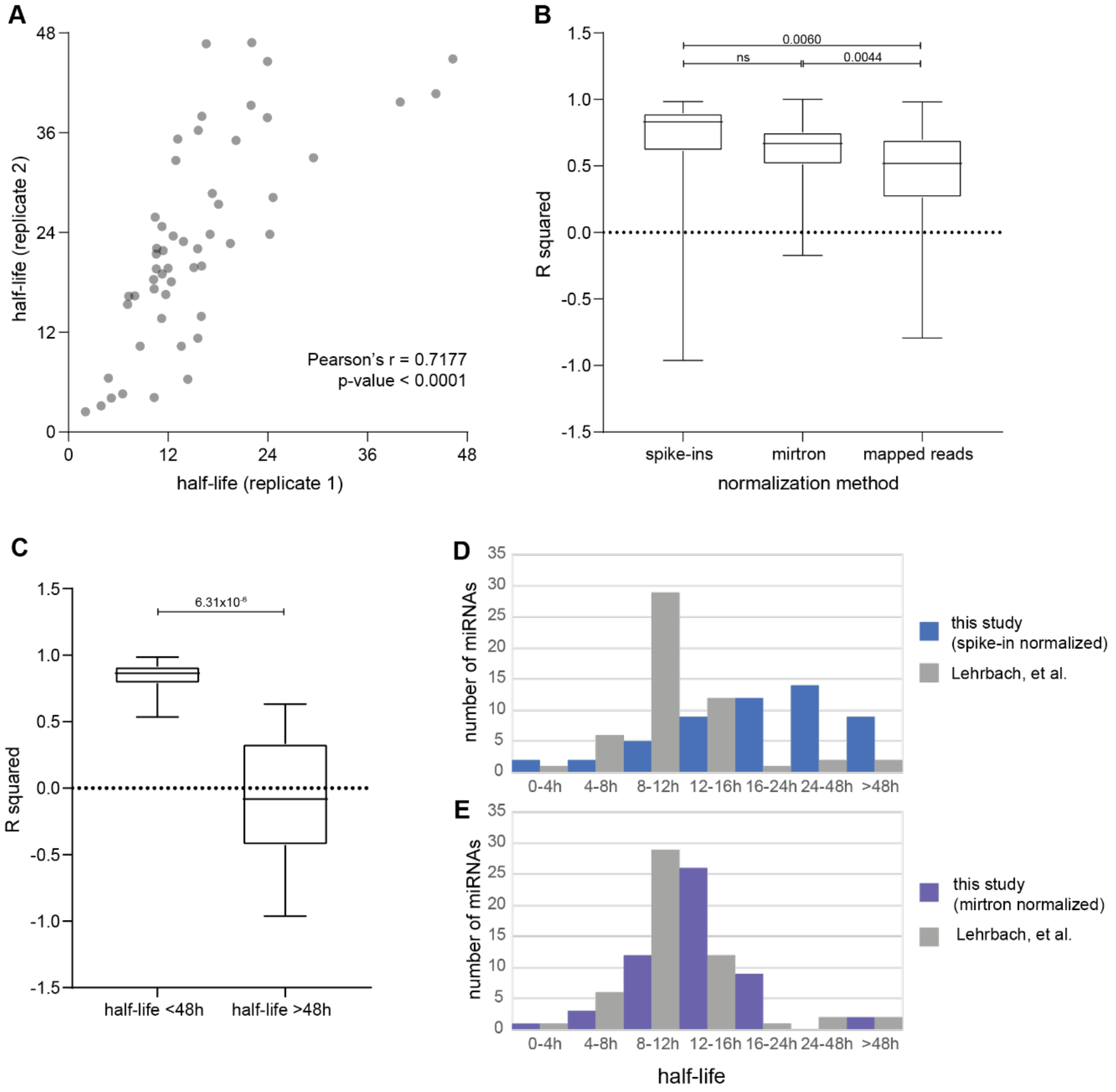


**Figure S10. Summary data from half-life calculations.** Only miRNAs with >50 RPM at 0h in both replicates are included in analysis, and annotated mirtrons are excluded. (A) Correlation between half-lives calculated using data from only one biological replicate. Half-lives greater than 48h are excluded due to low confidence (low R squared values). Pearson correlation coefficient and p-value of Pearson correlation test are shown. (B) Distribution of R squared values from fitting *pash-1(ts)* time course data to exponential decay curves (with data from both replicates), using three different normalization methods. Spike-in normalization performs best. One-way ANOVA test for significance was performed, followed by Tukey's multiple comparisons test (p-values shown). (C) Distribution of R squared values for half-lives above or below 48h. Half-lives within the time course of the experiment have a better fit. p-value from two-tailed heteroscedastic t-test shown. (D-E) Distribution of half-lives in a previous *pash-1(ts)* study in which a mirtron and siRNA were used for normalization, compared to half-lives calculated in the current study by two different normalization methods.

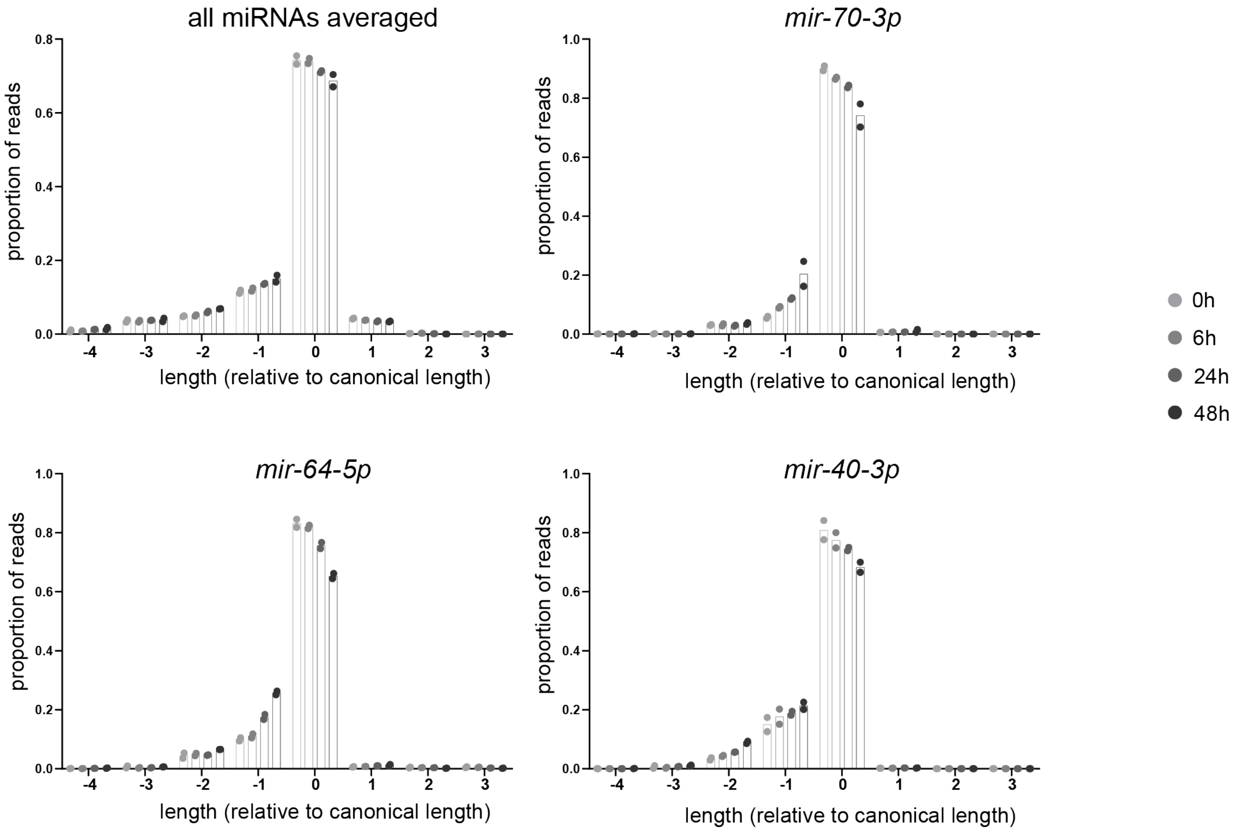


**Figure S11. Trimming of miRNAs over time course after PASH-1 inactivation.** Proportion of reads are plotted at each length, relative to the canonical length of the miRNA. In upper left panel, these proportions are averaged across all miRNAs that are >50 RPM in all time points.

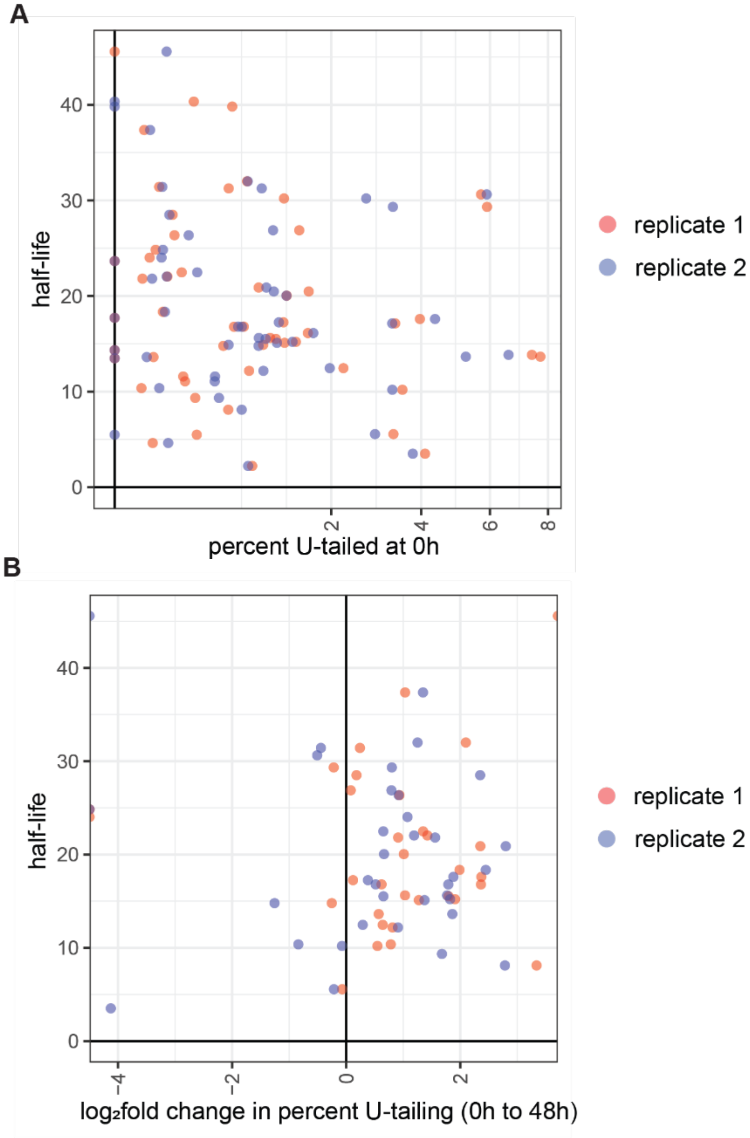


**Figure S12. Lack of correlation of half-life with mono-uridylation.** (A-B) Each dot represents one miRNA; each color is an independent biological replicate. Only miRNAs with half-lives <48h (and therefore of higher confidence) are shown. Pearson correlation test p-value > 0.05 for each replicate on each plot. (A) Correlation of half-life and percent mono-uridylation at 0h of *pash-1(ts)* time course. Only miRNAs with >50 RPM at the 0h time point are shown. (B) Correlation of half-life and change in mono-uridylation over time course, represented by $\log_2$fold change from 0h to 48h. miRNAs with >50 RPM in both the 0h and the 48h time point are shown.

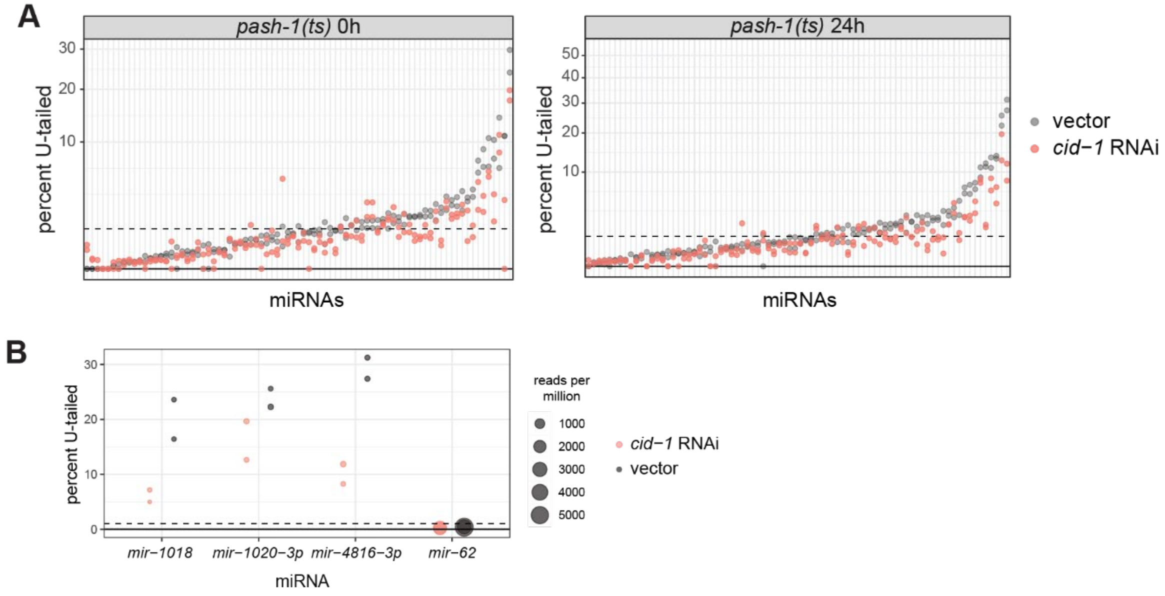


**Figure S13. Impact of untemplated uridylation on miRNA abundance and stability.** (A) Percent mono-uridylation in *pash-1(ts)* samples in empty vector or *cid-1* RNAi. All miRNAs with >50 RPM in each empty vector replicate of the indicated time point are shown. (B) Uridylation and abundance of mirtrons in *pash-1(ts)* samples 24h after upshift to restrictive temperature. All 3p-derived mirtrons with >25 RPM are shown. (Note a lower abundance cutoff is used - since modification levels are generally high - to enable visualization of all data points.)

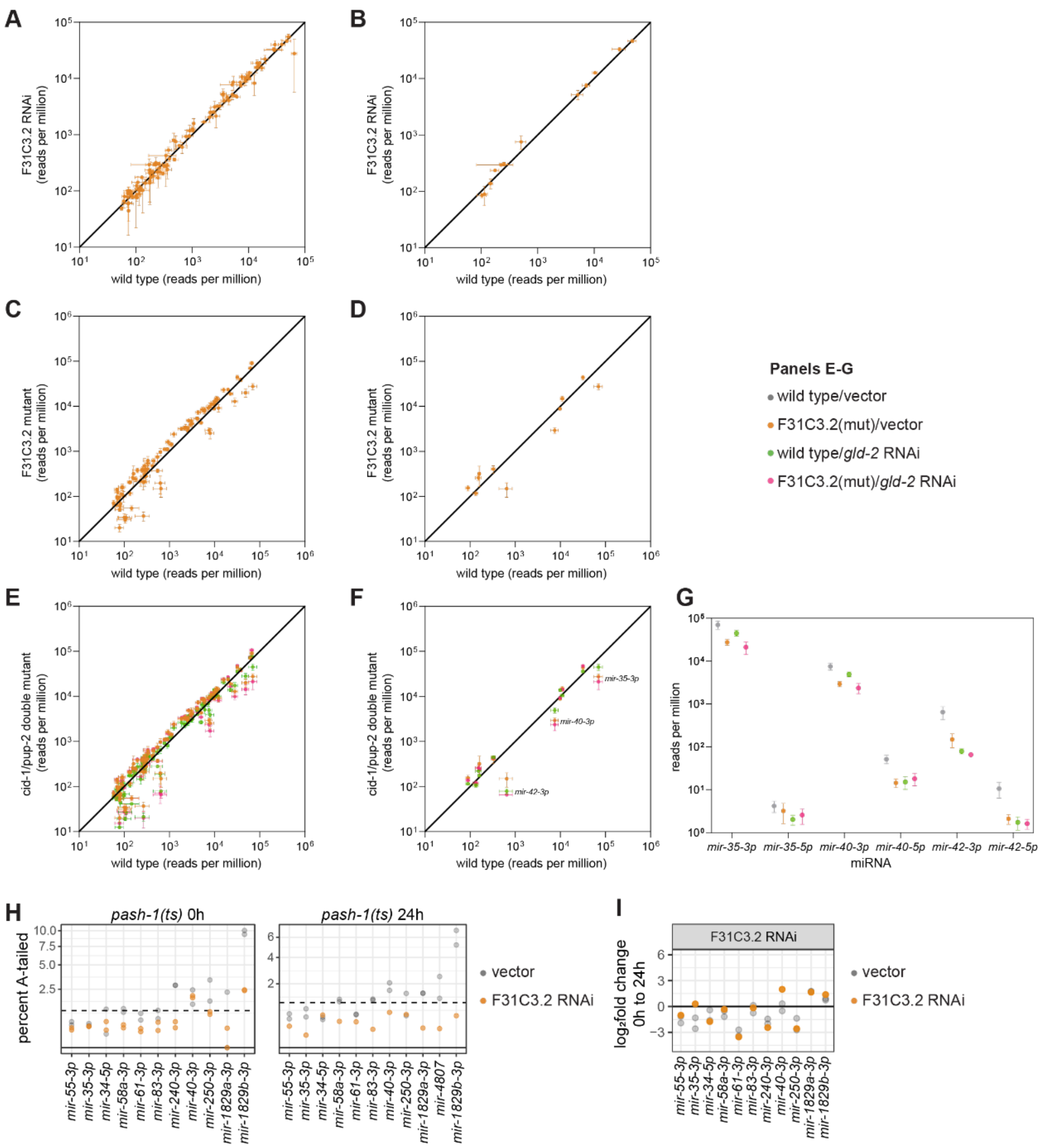


**Figure S14. Impact of untemplated adenylation on miRNA abundance and stability.** (A-B) Abundance of miRNAs in F31C3.2/*gldr-2* RNAi (two biological replicates) versus empty vector (three biological replicates). Mean and standard error are plotted. (A) All miRNAs >50 RPM in all empty vector replicates. (B) All miRNAs >50 RPM in all empty vector replicates and >1% average mono-adenylation across empty vector replicates. (C-F) Abundance of miRNAs in indicated genotype/RNAi versus wild type/empty vector. Mean and standard error of four biological replicates are shown. (C,E) All miRNAs >50 RPM in all wild type/empty vector replicates shown. (D,F) All miRNAs >50 RPM in all wild type/empty vector replicates and >1% average mono-adenylation across wild type/empty vector replicates are shown. (G) Abundance of guide (3p) and star (5p) strands of *mir-35-42* family members labeled in (F). (H) Percent mono-adenylation in empty vector or F31C3.2/*gldr-2* RNAi. A miRNA is shown if it is <50 RPM in both empty vector replicates of the indicated time point and the empty vector replicates average >1% mono-adenylation in wild type or either *pash-1(ts)* time point. (I) $Log_2$fold change from 0 to 24h after PASH-1 inactivation in empty vector (gray) or F31C3.2/ *gldr-2* RNAi. miRNAs are shown if they are >50 RPM in each empty vector of the 0h time point and the empty vector replicates average >1% mono-adenylation in wild type or either *pash-1(ts)* time point.

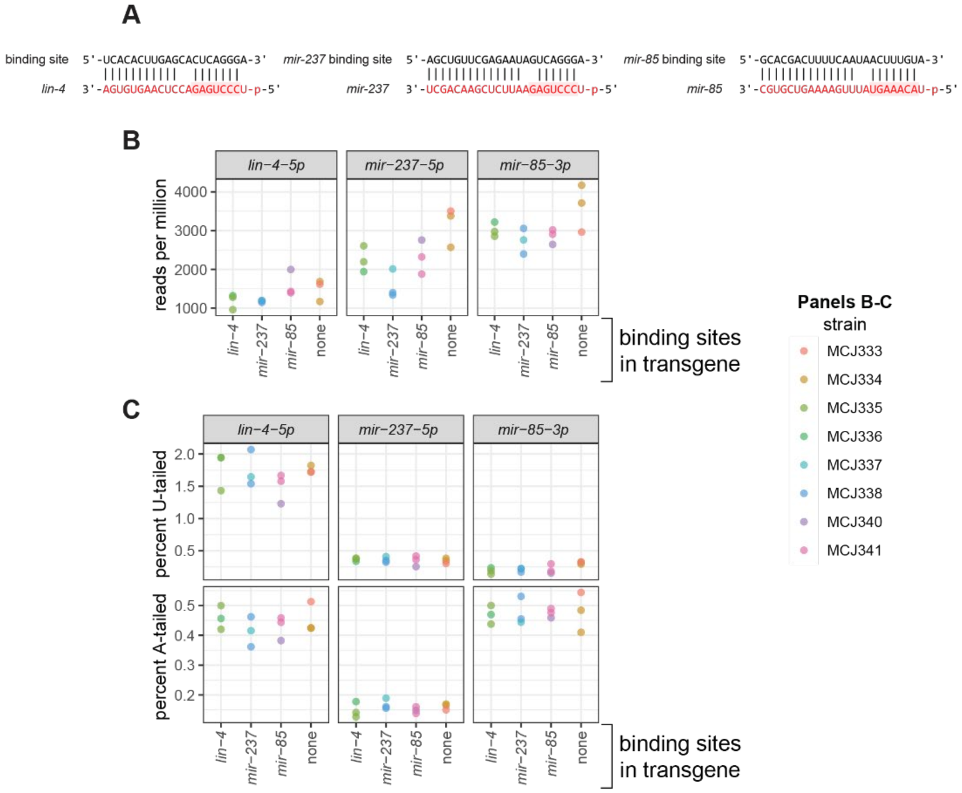


**D**

| | *lin-4* | *mir-237* | *mir-85* | *no site* |
|---|---|---|---|---|
| average molecules of miRNA (per 1ng total RNA) | 888,689.15 | 130,242.28 | 887,676.54 | |
| average molecules of mRNA trigger (per 1ng total RNA) | 352.22 | 270.33 | 2,576.00 | 3,219.67 |
| ratio (miRNA:mRNA) | 2,523.13 | 481.78 | 344.59 | |

**Figure S15. Standard over-expression transgenes unlikely to induce TDMD in *C. elegans*.** (A) Representative schematic of base-pairing at miRNA binding sites in over-expression transgenes. (B) Abundance of miRNAs complementary to expressed transgenes. (C) Percent mono-uridylation and mono-adenylation of miRNAs of interest. (B-C) Quantified miRNA is listed at top of plot. The x-axis labels denote which miRNA binding sites are contained in the expressed transgene. Plots are color coded according to strain. Note that two strains (two distinct transgenes) are examined for each miRNA. (D) Results of absolute quantification miRNA-Taqman using synthetic miRNAs to generate standard curve (top row), digital droplet PCR for absolute quantification of transgene mRNA (middle row), and ratio of these two species (bottom row).
